# Exploring the Structural and Functional role of α,β-unsaturated Ketoesters as Anti-Staphylococcal Agents Targeting Glutathione Peroxidase

**DOI:** 10.64898/2026.06.16.732695

**Authors:** Sushobhan Maji, Sayoni Dam, Akanksha Kumari, Himani Sharma, Naveen Sharma, Nirmal K. Rana, Asmita Samadder, Sudipta Bhattacharyya

## Abstract

Multiple-drug resistant (MDR) *Staphylococcus aureus* strains (like methicillin-resistant *S. aureus* or MRSA) uses an arsenal of antioxidant enzymes to mitigate host-induced oxidative stress. Among them the non-canonical Staphylococcal glutathione peroxidase (SaGpx) plays a crucial role in bacterial redox homeostasis by reducing peroxides via thioredoxin-dependent pathways. Thus, enabling oxidative stress mitigation during host infection. Despite its importance in *S. aureus*, its role in bacterial pathogenesis remains unexplored. This study aimed to elucidate the possible role of SaGpx in Staphylococcal virulence. First, we determined the high-resolution crystal structure of SaGpx (at 1.65 Å resolution) using X-ray crystallography. Guided by the catalytic cleft architecture of SaGpx, small-molecule based inhibitors were then rationally designed and synthesized. These inhibitors exhibited good binding affinity to SaGpx and complete enzymatic blockade. These inhibitors exhibited potent anti-*S. aureus* activity (MICs 6.25-31.25 μM) along with no cytotoxicity in L929 fibroblast wound-healing assays. Furthermore, the in vivo antibacterial ability of these inhibitors was evaluated using *S. aureus*-infected skin wound mouse model, where these compounds show potent antibacterial and wound healing ability supported by subsequent histological as well as immunohistochemical analysis. These findings suggest SaGpx as a possible virulence determinant in *S. aureus* and position these synthesized inhibitors as promising antivirulence therapeutics.

**Highlights:**

- The high-resolution crystal structure of Staphylococcal glutathione peroxidase is solved.
- Based on the SaGpx catalytic site, α,β-unsaturated ketoesters derivatives are synthesized.
- Synthesized α,β-unsaturated ketoesters derivatives inhibit SaGpx activity and binds the protein at μM range.
- Synthesized α,β-unsaturated ketoesters derivatives show in vitro antibacterial activity against *S. aureus* at low μM range.
- Synthesized α,β-unsaturated ketoesters derivatives show in vivo antibacterial and wound healing ability *S. aureus*-infected skin wound mouse model.

## Introduction

*Staphylococcus aureus* is a Gram-positive, non-motile, non-spore forming cocci shaped bacteria. Around 30% of the human population is colonized with *S. aureus*. Skin and anterior nares are the sites in human beings where this pathogen generally colonizes, making this commensal a part of the normal microbiota in the human host. It manifests pathogenicity particularly in immunocompromised hosts, causing a spectrum of diseases from mild skin and soft tissue infections to life-threatening conditions like septicemia and endocarditis ^1^. In addition, the emergence of multiple-drug resistant strains (particularly methicillin-resistant *S. aureus* or MRSA) worldwide complicated the outcome of treating this pathogen ^2^. The current treatment relies primarily on vancomycin. However, the emergence of vancomycin-intermediate (VISA) and resistant (VRSA) strains, poor clinical outcomes, and increased nephrotoxicity associated with high-dose therapy with vancomycin underscores the urgent need for novel therapeutic leads ^3,4^.

During infection *Staphylococcus aureus* faces extreme oxidative stress and depends on antioxidative defense mechanism for survival in hostile intracellular microenvironment of the host ^5^. The antioxidative defense mechanism is composed of a plethora of antioxidant enzymes including catalase, superoxide dismutase, and peroxidases ^6^. Recently, we have characterized one glutathione peroxidase (Gpx) homologs from *Staphylococcus aureus*, SaGpx (Uniprot Id: Q2FYZ0), an important member of the pathogens’ robust antioxidative defence system, which shows characteristic thiol dependent peroxidase activity ^7^. SaGpx reduces substrate hydroperoxides into water and less harmful alcohols, thereby maintains the cellular redox balance. Gpx is the terminal component of a specific enzyme cascade that works with the help of other cognate electron donors, generally glutathione (GSH) and glutathione reductase (GR) in case of classical Gpxs. *Staphylococcus aureus* neither possesses the biosynthetic machinery for GSH, the immediate reducing partner of Gpx, nor GR ^8^. Instead, this newly characterized SaGpx uses Staphylococcal thioredoxin 1 (SaTrx1) and Staphylococcal thioredoxin reductase (SaTR) as the reducing partner in the enzyme cascade, which ultimately draws electrons from NADPH. This non-canonical Staphylococcal glutathione peroxidase reduces substrate hydroperoxide with an apparent catalytic efficiency (with cumene hydroperoxide) of 2.1x10^4^ M^-1^·s^-1^, making it one of the key hydroperoxide stress mitigators in *S. aureus*. Site-directed mutagenesis studies indicate the involvement of catalytic tetrad residues composed of C36, Q70, W124 and N125 in catalysis. SaGpx follows a similar catalytic mechanism to that of atypical 2-Cys peroxiredoxin, where C36 acts as peroxidative cysteine (C_P_), and C82 acts as resolving cysteine (C_R_). Whereas the non-cysteine residues like Q70, W124 and N125 maintain the C36 in nucleophilic state (C_36_-S^-^) and are also responsible for substrate stabilization and efficient peroxide detoxification in SaGpx. The catalysis starts with the oxidation of the thiolate anion that results in the formation of sulfenic acid (S-OH). This intermediate state is then resolved by C_R_, resulting in the formation of a stable disulfide bond (S-S) between C_P_ and C_R_. This oxidized form of SaGpx is then recycled by SaTrx1 **(Scheme 1)** ^7^.

**Scheme 1.**
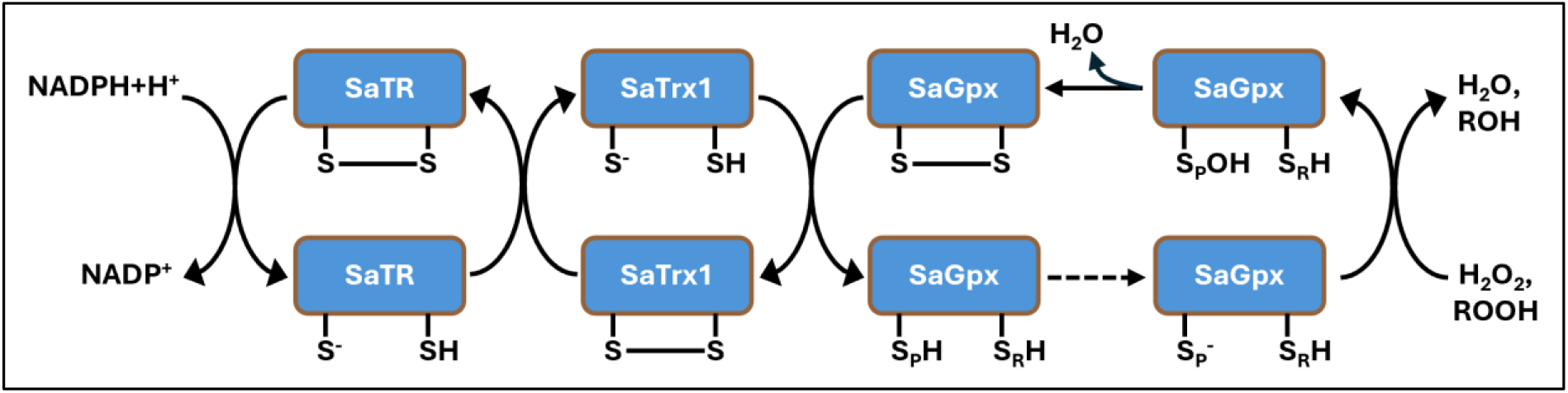
Electron relay in the catalysis of Staphylococcal glutathione peroxidase (SaGpx). The electron flows from NADPH+H^+^ via Staphylococcal thioredoxin reductase (SaTR), Staphylococcal thioredoxin 1 (SaTrx1) to reduce peroxides into their corresponding alcohols or water.

Studies in *Streptococcus pyrogens* highlight the critical role of Gpx in bacterial pathogenesis. The loss of GSH peroxidase activity (in Gpx-deficient strain) exhibits reduced ability to cause lethal systemic infection, underscoring the enzymes’ contribution to pathogen’s virulence ^9^. Previous studies also show that, among other antioxidant enzymes, the expression of glutathione peroxidase in *S. aureus* is also upregulated immediately after phagocytosis ^10^. Based on the literature pointing towards Gpx’s possible role in bacterial virulence, coupled with the crucial role of SaGpx in maintaining bacterial redox balance, motivated us to investigate the SaGpx’s plausible role in Staphylococcal virulence.

Herein our study, the plausible role of SaGpx in Staphylococcal virulence was probed by designing small molecule-based inhibitors targeting the active site thiolate anion of SaGpx. These inhibitors were designed based on the active site catalytic cleft of SaGpx. Kinetic analysis revealed that the SaGpx C36S mutant lost catalytic activity completely ^7^. Therefore, we hypothesized that an inhibitor that may modify the active site thiolate of SaGpx may also abolish its catalytic activity and may thereby compromise the survival of the bacteria under host-imposed oxidative stress and therefore, may also reduce the virulence of the bacteria. As a result, the host immune system will be able to clear the pathogen from the body.

α,β-Unsaturated carbonyl compounds are important pharmacophores because of their ability to undergo Michael addition with biological nucleophiles such as thiols, which makes them useful in many medicinal applications ^11^. Natural products like chalcones, curcumin, and zerumbone are well-known examples that contain this structural feature ^12^. The conjugated double bond present in these molecules acts as a Michael acceptor, contributing to their biological activity. Molecules containing this structural motif are known to exhibit a broad spectrum of biological activities, including anti-inflammatory ^13^, antioxidant ^14^, antiviral, antibacterial, and antifungal agents ^15^. Due to these biological activities, several molecules based on α,β-unsaturated carbonyl systems have been explored as potential drug candidates and are used in the treatment of diseases such as cancer ^11^ and multiple sclerosis ^16^. Apart from their medicinal relevance, these compounds are widely employed as useful synthons in organic synthesis for constructing a variety of functional molecules through carbon–carbon and carbon–heteroatom bond formation ^17^.

In this study, with our approach to develop small molecule-based therapeutic leads against *S. aureus* targeting Staphylococcal glutathione peroxidase, a series of small molecule derivatives having α,β-unsaturated carbonyl systems were synthesized. The synthesis of these derivatives was based on the high-resolution crystal structure of SaGpx (at 1.65 Å resolution) obtained using X-ray diffraction crystallography. These derivatives have an ester functionality at the β-carbon and benzoyl carbonyl on the other side attached with different electron-withdrawing groups. The inhibitory activity of these synthesized derivatives against SaGpx was assessed through the determination of respective binding affinity (K_d_) values, followed by an enzyme inhibition assay. Subsequently, the antibacterial activity of these compounds was also assessed in vitro. Finally, the in vivo antibacterial activity and wound healing efficacy of these compounds were assessed in *S. aureus*-infected skin wound mouse model.

## Results and Discussion

### 1. Overall Structure of SaGpx and Active Site

To understand the structural basis of SaGpx function and to provide a framework for inhibitor design, we first determined the crystal structure of SaGpx. Initially we tried to get the crystal structure of wild type SaGpx in reduced and oxidized form, however after several trial and optimization we could not get diffraction quality crystals. Hence, we then tried with the catalytically attenuated SaGpx C36S mutant, which crystallizes **(Figure S2A)** and provides high-resolution three-dimensional structural information at 1.65 Å **(Figure S2B)**.

The SaGpx C36S mutant adopts a globular conformation with overall dimensions of approximately 31.2 Å x 36.8 Å x 38.2 Å. The protein exhibits the characteristic glutathione peroxidase fold, primarily composed of a centrally aligned twisted β-sheets formed by parallel (β3-β4-β5) and antiparallel (β3-β6-β7) strands. This core is flanked by four main α-helices, with helices α3, α4, and α7 situated on one side of the β-sheet, and helix α5 on the opposite side **(Figure 1A)**. The reduced conformational state of this structure is ensured by well resolved electron densities for the free sulfhydryl groups of C64 and C82, as well as the absence of any intramolecular disulfide bonds. The active site region is composed of residues C36, Q70, W124, and N125, all these collectively form the catalytic tetrad **(Figure 1B,C)**. Also the crystal structure shows electron density for four sulfate ions (came from crystallization solution) and six hydrogen peroxide molecules, which are most likely generated by radiation during data collection, as no hydrogen peroxide was added during crystallization or diffraction experiments. Table S1 summarizes crystal diffraction data collection, processing, and refinement statistics.

**Figure 1.**
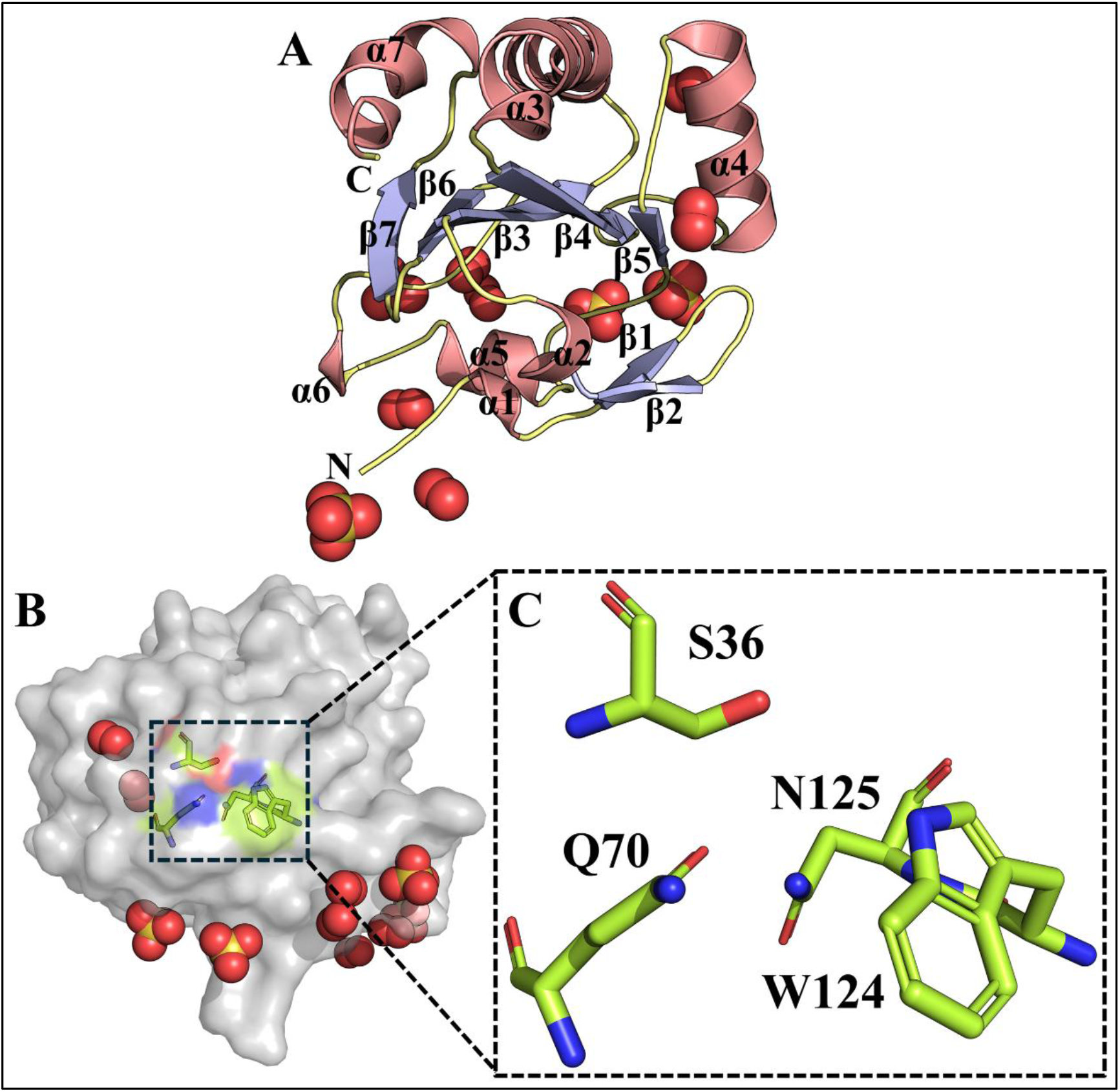
Overall structure of SaGpx C36S mutant. (A) Crystal structure as cartoon representation revealing the canonical Gpx-fold consisting of central β-sheet (blue) surrounded by α-helices (pink) with N-terminal secondary elements. (B) Surface view highlighting the catalytic tetrad (yellow) within the active site cleft. (C) Enlarged view of the catalytic tetrad (as the structure solved from SaGpx C36S mutant variant, S36 corresponds to the C36 in the native protein).

### 2. Ketoester Derivatives Exhibits High Binding Affinity and Potent Enzymatic Inhibition Against SaGpx

The structural elucidation of the SaGpx active site provides crucial insights into its catalytic mechanism, which hinges on the redox cycling of critical cysteine residues ^7^. Kinetic and site-directed mutagenesis data have established that SaGpx’s catalysis is driven by a highly reactive peroxidative cysteine (C36). The catalytic mechanism of SaGpx starts with the nucleophilic attack of the peroxidative cysteine thiolate (C_P_-S^-^) to the polarized peroxyl bond of the substrate hydroperoxide **(Scheme 1)**. This attack generates the reaction intermediate, sulfenic acid (R-SOH), which is subsequently resolved by resolving cysteine (C_R_). The disulfide bond thus formed between the C_P_ and C_R_ is recycled back to thiol form with the help of Staphylococcal thioredoxin 1 (SaTrx1). Mutating the peroxidative cysteine to serine completely attenuates SaGpx’s catalytic activity ^7^.

Because this thiolate anion is both highly reactive and absolutely essential for the initial step of the substrate peroxide reduction, it presents an ideal therapeutic target. We hypothesized that an inhibitor which may modify the active site thiolate of SaGpx may also abolish its catalytic activity and would arrest catalytic turnover. Therefore, the design of small-molecule-based SaGpx inhibitors was based on their electrophilic α,β-unsaturated carbonyl system, predicted to undergo Michael addition with the nucleophilic thiolate (C_36_-S^-^) generated at the enzyme active site. Initial screening confirmed the thiol-modifying capacity of compound C1, with a K_d_ value of 116.77 µM. To enhance the potency of this molecular scaffold, derivatives were synthesized by introducing electron-withdrawing groups on the aromatic ring, yielding markedly lower K_d_ values **(Table 1, Figure S6)**.

**Table 1.**
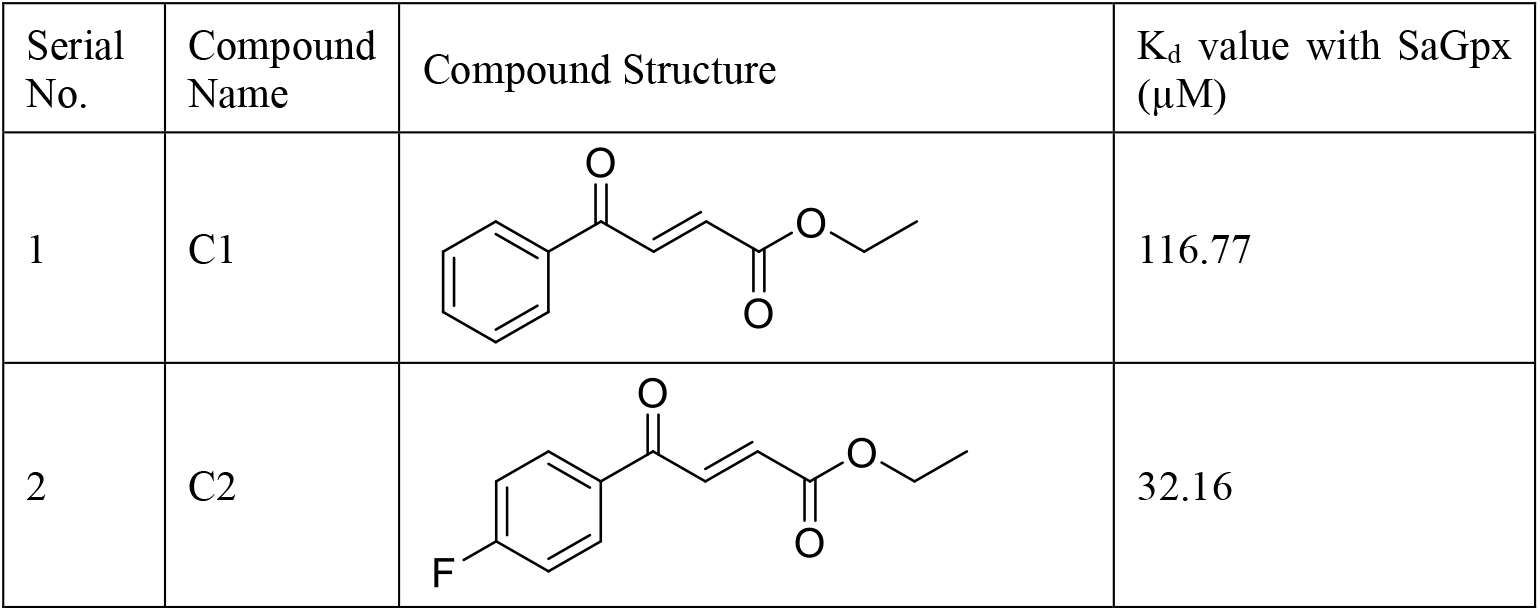
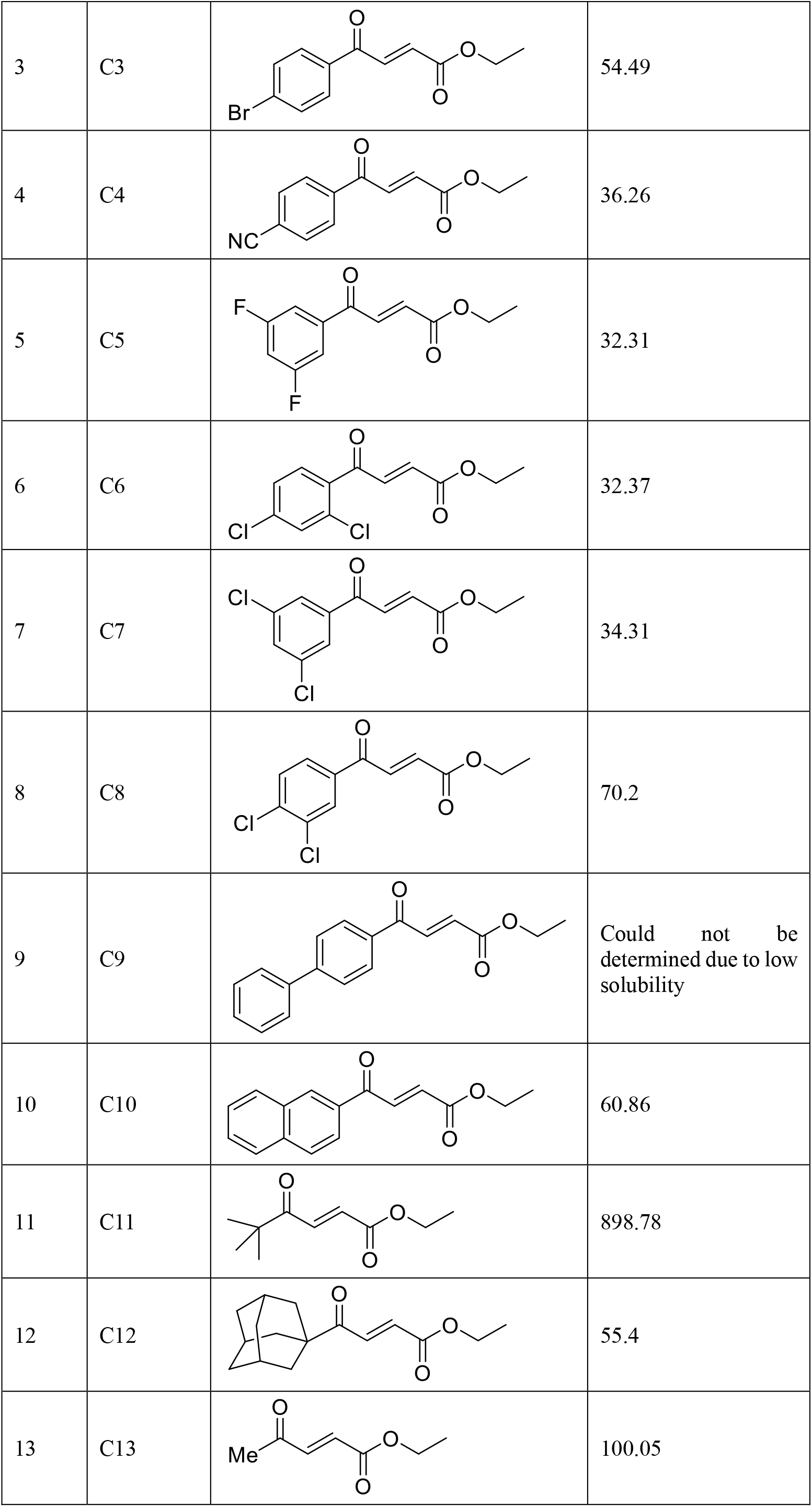
Binding Affinities (K_d_ Values) of the Synthesized ketoester Derivatives Against SaGpx.

| Serial No. | Compound Name | Compound Structure | $K_d$ value with SaGpx (μM) |
| --- | --- | --- | --- |
| 1 | C1 |  | 116.77 |
| 2 | C2 |  | 32.16 |
| 3 | C3 |  | 54.49 |
| 4 | C4 |  | 36.26 |
| 5 | C5 |  | 32.31 |
| 6 | C6 |  | 32.37 |
| 7 | C7 |  | 34.31 |
| 8 | C8 |  | 70.2 |
| 9 | C9 |  | Could not be determined due to low solubility |
| 10 | C10 |  | 60.86 |
| 11 | C11 |  | 898.78 |
| 12 | C12 |  | 55.4 |
| 13 | C13 |  | 100.05 |

The synthesized ketoester derivatives have an α,β-unsaturated double bond with an ester group on one side and a benzoyl carbonyl on the other. These groups pull the electrons from the double bond and create a partial positive charge at the β-carbon **(Figure S3)**. The ester group withdraws electrons only moderately because its alkoxy group donates electron density by resonance. In contrast, the benzoyl group withdraws electrons much more strongly, as both the carbonyl and the attached phenyl ring pull the electrons. This makes the β-carbon more electrophilic and allows the thiolate anion generated at SaGpx C36 to attack it easily **(Figure S4)**. The addition of an electron-withdrawing group to the benzene ring, such as F, Br, etc., increases the overall electron-withdrawing capacity of the benzoyl group, which ultimately increases the electrophilicity of the β carbon of the double bond. This enhanced electrophilicity resulted in greater interaction with the nucleophilic center, which is reflected by the reduced K_d_ values of compounds bearing additional electron-withdrawing groups like compound C2 (K_d_ = 32.16 µM), C3 (K_d_ = 54.49 µM), C4 (K_d_ = 36.26 µM), C5 (K_d_ = 32.31 µM), C6 (K_d_ = 32.37 µM), and C7 (K_d_ = 34.31 µM) compared to C1 **(Table 1)**. Structure-activity relationship analysis revealed that the position of the electron-withdrawing group also influences the electron-withdrawing capacity. For example, the shifting of chlorine from ortho (C7) to meta (C8) position increased the K_d_ values from 34.31 µM to 70.2 µM. Along with electronic effects, the binding pose also influences how these small molecules interact with SaGpx. The active site of the SaGpx is surrounded by hydrophobic amino acid residues like W124 and F38 **(Figure S5)**. W124 is involved in maintaining the thiolate anion form of peroxidative cysteine (C36). F38 is surface exposed and based on its three-dimensional disposition, it is likely that F38 may play a significant role in the binding of these small molecules through pi-pi stacking. Compound with extended aromatic systems (such as C10) or rigid, cage-like structures in case of adamantane-substituted ketoester derivative C12, show low K_d_ values, suggesting their efficient hydrophobic packing and subsequent binding and interaction in the hydrophobic environment near the SaGpx active site. In contrast, compounds like C11, where the benzene ring was substituted with a tert-butyl group (in order to check whether this aromatic group is critical in inhibitor’s efficient binding), show a significant increase (∼9 times) in K_d_ value as compared to C1. The more flexible tert-butyl group likely packs less effectively with the hydrophobic active site pocket compared to the rigid aromatic ring, resulting in weaker binding affinity and a higher K_d_ value. All these findings suggest the significance of the benzene ring in the proper binding of these small-molecule-based inhibitors within the shallow active site pocket of SaGpx. However, to know the exact mechanism of interaction of these small molecules, co-crystal structures of these compounds with SaGpx would be very helpful. The co-crystal structures would also be helpful in designing new derivatives with increased potency. Also, it would be imperative to study the interaction between the F38 mutant variant of SaGpx with these compounds to probe the role of F38 in these synthesized small-molecule binding.

These ketoester derivatives were synthesized to interact with the nucleophilic thiolate of Cys36 within the active site of SaGpx. The binding affinity assay was based on detecting the free thiol group of SaGpx in the presence or absence of these compounds. Upon titrating these compounds with SaGpx, the amount of available free thiol groups in the enzyme decreases. Following this binding affinity analysis, we seek to verify if these compounds directly interact with the Cys36 thiolate and render the enzyme catalytically inactive. To accomplish this, we evaluated their enzyme-inhibitory potential. We selected the two most promising compounds (C2 and C5, which demonstrated the lowest K_d_ values) and assessed their inhibitory effects against SaGpx using the FOX assay (as described in the Materials and Methods section). Both compounds potently inhibited SaGpx activity, blocking the electron transfer from the enzyme to the peroxide substrate **(Figure 2)**. Collectively, these binding and inhibition data suggest a possible covalent modification of the active site thiolate. However, co-crystal structures of these compounds with SaGpx remain necessary to provide definitive confirmation.

**Figure 2.**
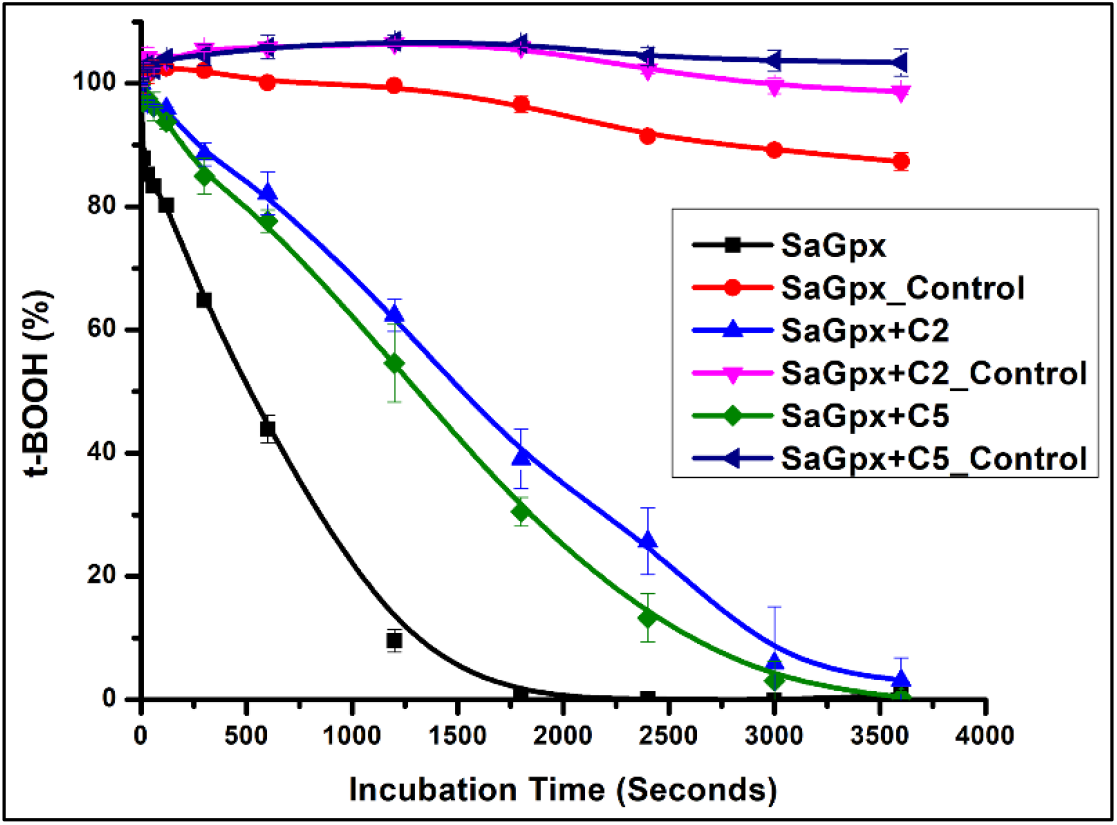
Enzymatic inhibition assessment of compound C2 and C5 and SaGpx using FOX assay. In case of SaGpx_Control, SaGpx+C2_Control and SaGpx+C2_Control samples, SaGpx was not added in the reaction mixture.

### 3. Ketoester Derivatives Shows Potent Antibacterial Activity Against *S. aureus*

After confirming the inhibitory potential of the synthesized derivatives against SaGpx, we next evaluated their broader antibacterial activity. First, we began with standardized agar well diffusion assays, a classical qualitative method where compounds are pipetted into wells punched into *S. aureus* seeded agar plates. Inhibition manifests as clear zones of growth suppression (zones of inhibition, ZOI) around the wells, proportional to the potency. Nearly all the ketoester derivatives exhibited antibacterial activity against *S. aureus* **(Figure 3, Figure S7)**. Most of these compounds showed antibacterial activity comparable to kanamycin, with compounds C3, C5, C6, C7, C8, and C12 showing better antibacterial potential than kanamycin. However, compounds C11 and C13 did not show any antibacterial activity.

**Figure 3.**
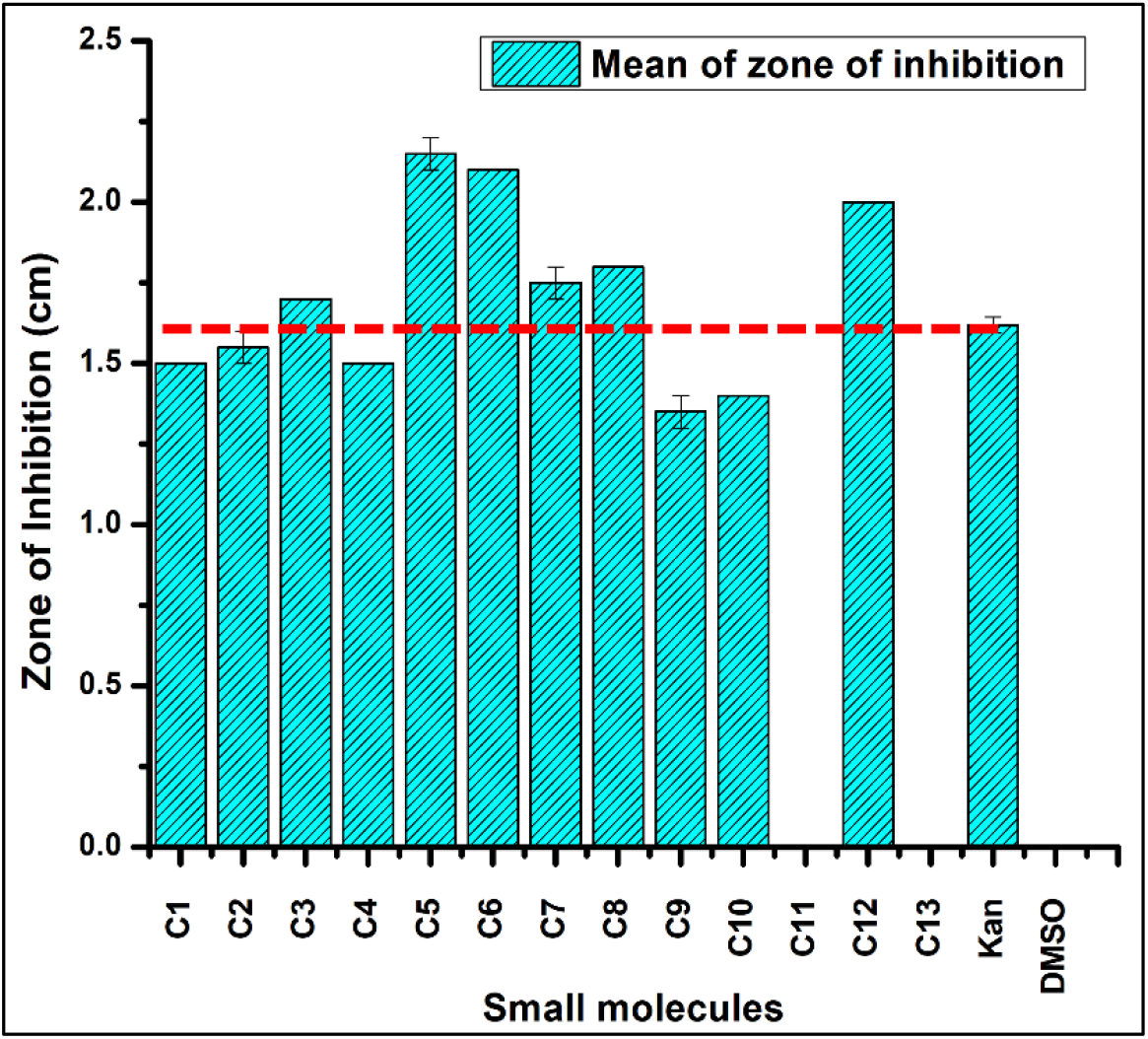
Graphical representation of the Zone of Inhibition of the tested inhibitors through the agar well diffusion assay.

Following the agar well diffusion assay results, the minimum inhibitory concentration (MIC) values of the compounds showing antibacterial activity against *S. aureus* were determined. These MIC values were calculated using standard broth microdilution methods. All the tested compounds exhibited MIC values in the range of 6.25 µM to 31.25 µM **(Table 2)**.

**Table 2.** MIC Values of the Synthesized Ketoester Derivatives Against *S. aureus*.

| Serial No. | Compound Name | MIC Values (μM) against *S. aureus* |
| --- | --- | --- |
| 1 | C1 | 15.6 |
| 2 | C2 | 15.6 |
| 3 | C3 | 7.8 |
| 4 | C4 | 31.25 |
| 5 | C5 | 12.5 |
| 6 | C6 | 6.25 |
| 7 | C7 | 7.8 |
| 8 | C8 | 7.8 |
| 9 | C9 | 6.25 |
| 10 | C10 | 6.25 |
| 11 | C12 | 31.25 |

Following confirmation of the synthesized ketoester derivatives’ in vitro antibacterial activity, their in vivo efficacy was further assessed using *S. aureus*-infected skin-wound mouse model. Prior to these studies, potential adverse effects of these compounds on mammalian cells were examined through an in vitro cell migration and scratch closure assay using L929 fibroblast cells. For these experiments, compounds C6 and C7 were selected based on their low MIC values (6.25 µM and 7.8 µM, respectively) and solubility profiles.

### 4. Ketoester Derivatives Facilitate L929 Fibroblast Migration and Wound Healing

The effect of the ketoester derivatives on fibroblast migration was evaluated using an in vitro scratch closure assay in L929 cells. Both the compounds, C6 and C7, exerted no adverse effect on the cells. Neither was the morphology of these cells affected, nor was there inhibition in cellular migration over the 24 h. In both compounds at applied conditions, an overall similar rate of scratch healing was observed compared to the control group. For compound C6, the scratch closure rate was less than that of the control group at 2 h, followed by a marked increase over the next 4 h; however, by 24 h, the wound closure remained only 2.57± 0.157% lower than the control group. In contrast, compound C7 showed significantly higher scratch closure rate than control group at 2 and 6 h and achieved an almost comparable scratch closure rate at 24 h being only 0.156±0.538 % lower than the control group **(Figure 4A-4B)**.

**Figure 4.**
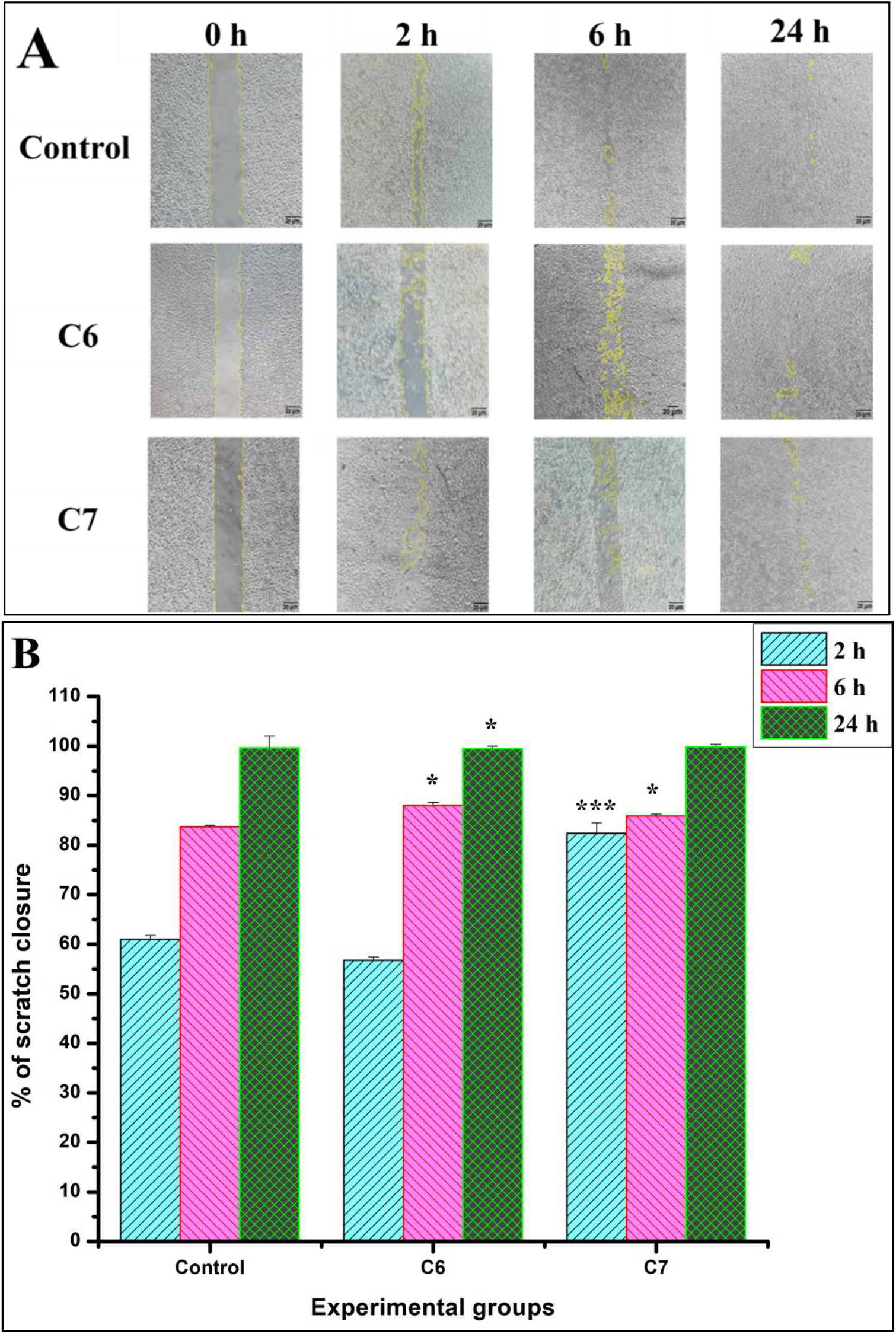
Cell migration and scratch closure analysis. (A) Migration of L929 cells into the scratched region demonstrating progressive wound closure. Yellow dotted lines indicate the border of the scratched area. (B) Graphical representation of scratch closure (%). *p<0.05, ***p<0.001 vs. Control, were considered significant for Student’s t-test.

Migration of cells in the scratched area triggered by different immunological factors is a natural phenomenon that results in closure of the scratched or wounded area. Both tested compounds exhibited pro-migratory effect on L929 fibroblasts. By simultaneously supporting the fibroblasts’ migration and scratch healing, these ketoester derivatives demonstrate dual functionality that is advantageous for wound healing applications, which are further evaluated in the subsequent mouse model experiment.

### 5. Ketoester Derivatives Accelerate Wound Healing and Reduce Inflammation in *S. aureus* Infected Skin-wound Mouse Model

#### 5.1. Macroscopic Images of Wound Healing

Macroscopic images of all experimental groups were captured from a fixed distance immediately after wounding, one day after inoculation, and on days 1, 7 and 14, to compare as well as assessing the wound healing progression. Macroscopic examination **(Figure 5)** shows that, 24 h after the introduction of the bacterial inoculum in mice, the wound area swelled significantly along with redness and appearance of yellowish pus-like exudate, indicative of the inflammatory phase of wound healing. In contrast, the non-infected “only wound” group showed very mild inflammatory signs and showed early signs of recovery. Day 1 refers to the first day after application of the inhibitor.

**Figure 5.**
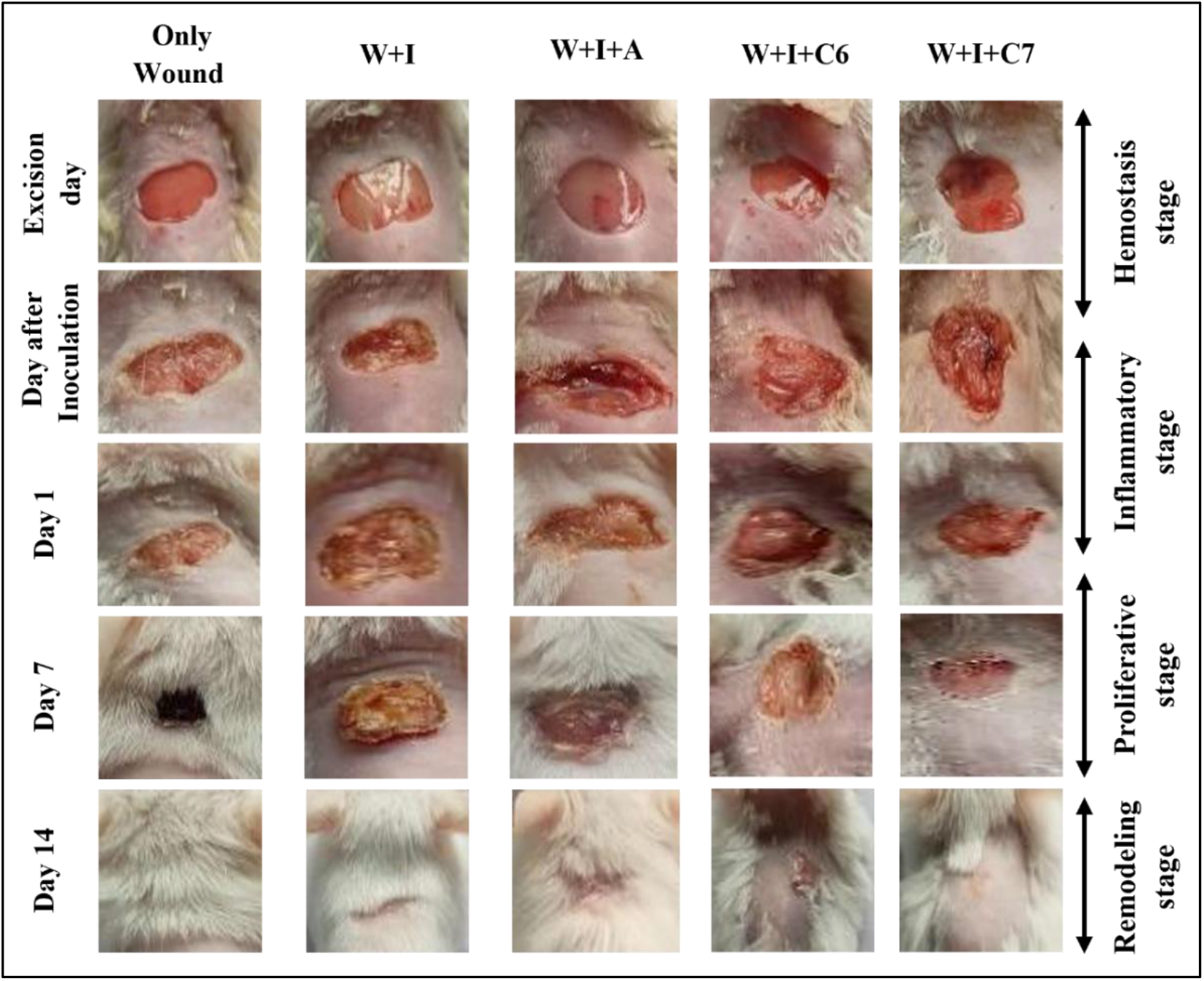
Macroscopic images of wound area in mice across all experimental groups. W, I, A, C6 and C7 represent wound, infection, antibiotic, compound C6 and compound C7 respectively.

On day 1, wound swelling and exudation were reduced more prominently in the “compound C7-treated group (W+I+C7)” compared with the “infected untreated group (W+I)” group and the “antibiotic-treated group (W+I+A)”. In the “compound C6-treated group (W+I+C6)”, swelling persisted, although the wound surface appeared drier superficially. In the “only wound” group, scab formation had already begun by day 1. By day 7, the wound area in the “only wound” group had retracted with minimal residual scab, whereas in the W+I group, the wound area remained larger with a prominent scab, consistent with the delayed transition into the proliferative phase. In the W+I+C6 group, swelling gradually decreased from day 1 but was still evident, and scab formation was not yet complete. In contrast, W+I+C7 group, the scab had detached from the wound area by day 7, indicating faster wound contraction and recovery compared to W+I+C6 group and other experimental groups, including W+I+A group. By day 14, scab detachment was complete in the “only wound” and W+I+A group, accompanied by dense hair regrowth over the healed area. But in case of W+I, i.e., only infected group, though scab detached but in the wounded area, signs of infection still persisted which remain hindered under newly grown fur, and the same instance also experienced in other mice of this experimental group. In both the W+I+C6 and W+I+C7 groups, fur regrowth was relatively scant compared to other groups. Overall, the macroscopic observations suggest that both compounds C6 and C7 possess antimicrobial activity coupled with wound-healing potential. Furthermore, compound C7 demonstrates superior efficacy to compound C6 in this infected wound model.

#### 5.2. Percentage of Wound Area Contraction

Skin is a rapidly growing tissue and represents the first line of defense against microbial invasion. Skin also harbors a diverse commensal microbiota. When skin wounds become contaminated with pathogenic microbes such as *S. aureus*, wound contraction is delayed, leading to prolonged infection ^18^. Wound repair is a complex, multistep process in which different immune factors coordinate to heal the wound. Wound closure involves progressive contraction, during which the wound edges are drawn together through the migration and interaction of fibroblasts, keratinocytes, and matrix components. Fibroblast cells migrate to close the wound area, keratinocytes become flattened and dead keratinocytes accumulate to make stratum corneum in order to restore the normal structure of skin ^19^.

In the present study, 24 h after bacterial infection, wound area increased by 17.94±0.29%, 28.020±7.29%, 26.33±3.71%, 14.98±3.75%, 29.25±4.69% in only wound, W+I, W+I+A, W+I+C6, and W+I+C7 groups respectively, relative to the excision day, consistent with infection-associated swelling. With the formula mentioned earlier, wound contraction rates were calculated with respect to wound area measured 24 h after inoculation. As summarized in Table 3, the wound closure rate of “only wound” group was much higher than W+I group. However, on day 1, wound closure rates in the C6 and C7 treated groups were significantly higher than both the earlier groups. On day 7 wound contraction rate reached 75.48 ± 2.36% and 65.88±2.895% in C6 and C7 treated groups, respectively, compared with 8.9±0.988% in W+I, clearly indicating the wound healing potential of selected compounds. Moreover, both compounds (C6 and C7) applied groups also showed more efficacy in terms of wound contraction rate than antibiotic treated groups and by 14th day wound healed completely **(Figure 6)**.

**Table 3:** Percentage (%) of wound closure (Mean ± standard error) of all experimental groups.

| Day | Only wound | W+I | W+I+A | W+I+C6 | W+I+C7 |
| --- | --- | --- | --- | --- | --- |
| Day after inoculation day | 0 % | 0% | 0% | 0% | 0% |
| Day 1 | 12.45<br>$\pm 1.629\%$ | 3.36<br>$\pm 0.25\%$ | 4.896<br>$\pm 0.353\%$ | 16.235<br>$\pm 1.56\%$ | 15.98<br>$\pm 2.358\%$ |
| Day 7 | 70.29<br>$\pm 1.745\%$ | 8.9<br>$\pm 0.988\%$ | 40.698<br>$\pm 3.356\%$ | 75.48<br>$\pm 2.36\%$ | 65.88<br>$\pm 2.895\%$ |
| Day 14 | 99.98$\pm 0.98$ | 86.9$\pm 1.38$ | 98.50$\pm 4.07$ | 99.69$\pm 1.823$ | 99.568$\pm 3.45$ |

**Figure 6:**
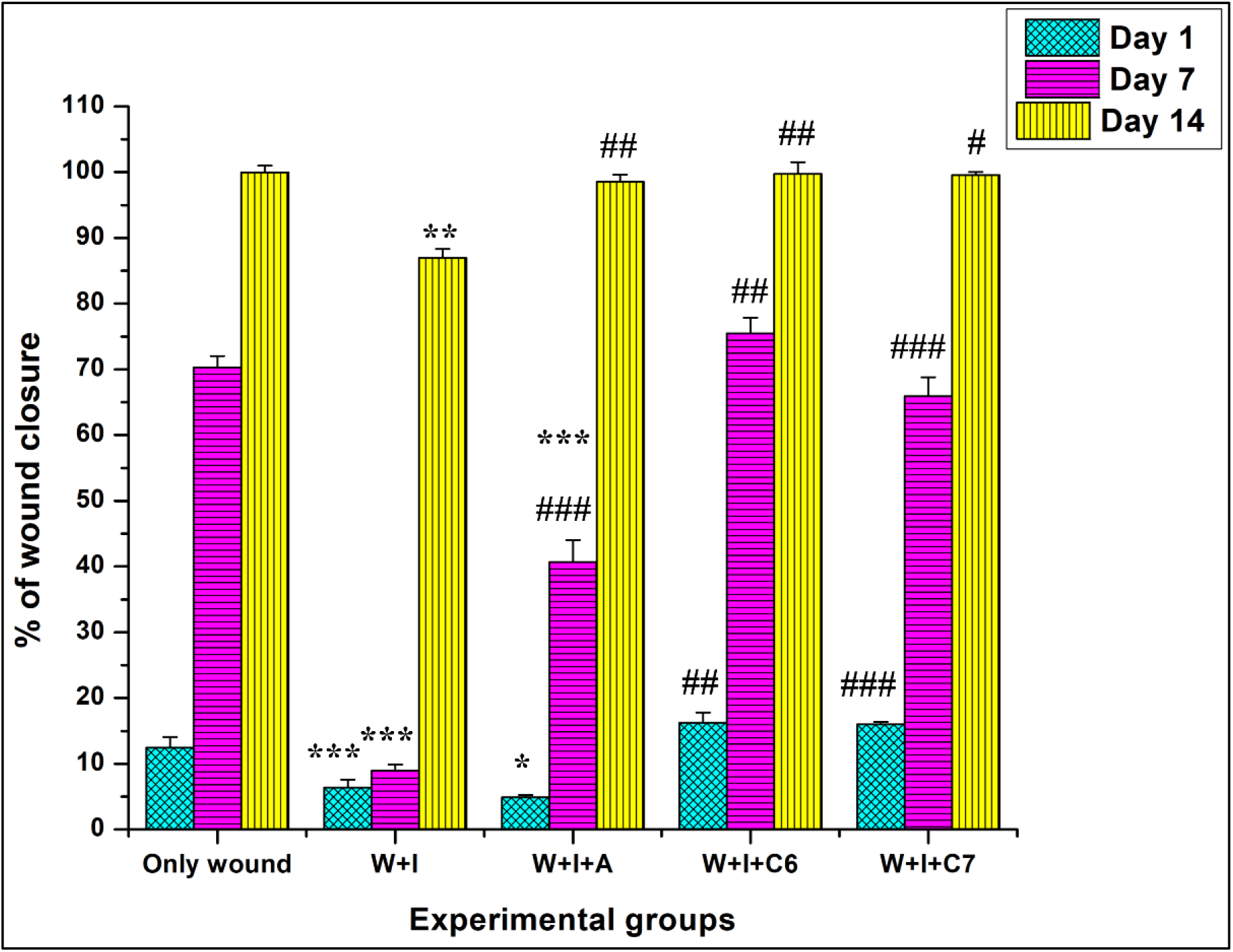
Graphical representation of the percentage of wound retraction, indicating the extent of wound healing across all experimental groups. *p<0.05, **p<0.01, ***p<0.001 vs. Only wound and ^#^p<0.05, ^##^p<0.01, ^###^p<0.001 vs W+I, were considered significant for Student’s t-test. W, I, A, C6, and C7 represent wound, infection, antibiotic, compound C6 and compound C7 respectively.

#### 5.3. Histopathological Findings

Histopathology serves as the gold standard for understanding and comparing changes in tissue architecture, paving the way for understanding the action of candidate compounds on specific organs. Skin wound healing is distinctively segregated into four main phases: haemostasis, inflammation, proliferation and remodeling. All these phases have distinct histological features, which explains sequential changes in wounded tissues ^20^.

As mentioned earlier, day 1 corresponds to 24 h after application of compounds. On day 1, the “no wound” group showed normal epidermal thickness, minimal tissue granularity and an intact collagen layer **(Figure 7)**. In the “only wound” group, epidermal hyperplasia and increased tissue granularity due to the deployment of neutrophils, macrophages, a typical feature of inflammatory phase, were evident but less pronounced than in the *S. aureus* infected group. In compound C6 and C7 treated groups, shearing and abnormal thinning of epidermis was observed, however tissue granulation and loosening of collagen tissue were less than other three groups, suggesting attenuation of inflammation. In the C7-treated group, flattened keratinocytes started depositing on the epidermal layer, marking the initiation of the cornification process. Overall, lower tissue granularity suggests decreased inflammation in the compound treated groups.

**Figure 7:**
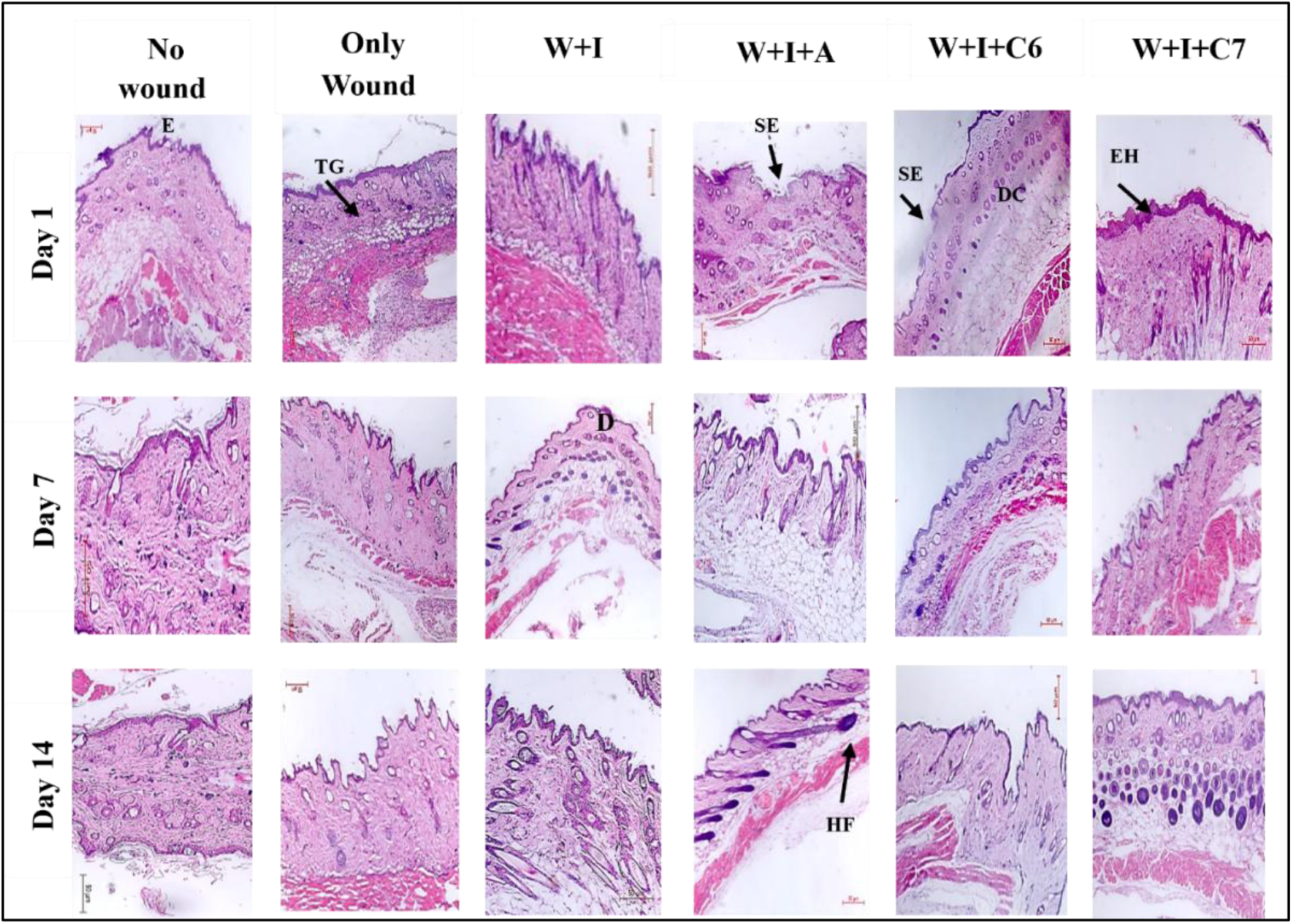
Hematoxylin-Eosin stained histological images of skin tissue of all experimental groups. Scale bar is 50 µm. All images are 10X magnified. Abbreviations: E: epidermis; D: dermis; TG: tissue granulation; SE: sheared epidermis; DC: depleted epidermis; EH: Epidermal hyperplasia; HF: hair follicles.

By day 7, the marked increase in epidermal layer thickness, a common immune response elicited due to injury, was decreased in the C7 treated group, and the shearing of the epidermal layer, which was observed in antibiotic treated and C6 treated group, had largely resolved. In both compound C6 and C7 treated groups, migration of keratinocyte, appearance of immature hair follicles and fibrous tissue were seen, and the collagen layer started attaining its normal thickness, indicating the onset of proliferative phase. By day 14, the collagen layer appeared more intact, epidermal layer became normal, the appearance of melanocytes and hair follicles become more abundant, although growth and number of hair follicles were more plentiful in “no wound”, W+I and antibiotic-treated groups. In C6 and C7 treated groups, formation of sebaceous gland and new blood vessels were visible, corroborating the macroscopic image of this group at this time point. Stratum corneum also appeared in these groups. Collectively, these histological findings support the wound healing efficacy of the selected compounds **(Figure 7)**.

#### 5.4. Immunohistochemical Analysis

Nuclear factor kappa B (NF-κB) is a pivotal transcription factor that orchestrates pro-inflammatory cytokine production (e.g., TNF-α, IL-6) in response to bacterial pathogen and oxidative stress like in *S. aureus*. Immunohistochemistry (IHC) provides spatial, semi-quantitative assessment of NF-κB nuclear translocation in tissue sections, making it ideal for evaluating inflammatory attenuation. IHC analysis revealed persistently elevated NF-κB expression in the W+I group across all the observation days (days 1, 7, and 14) with only gradual decline over time, which is characteristic of unresolved *S*.*aureus*-driven inflammation. In contrast, NF-κB levels in the C6 and C7 treated groups were markedly reduced by day 7 compared to day 1, indicating that these compounds modulated the immune response and attenuated inflammation. Notably, NF-κB expression levels on days 7 and 14 were also significantly lower in both compound C6 and C7 treated groups than antibiotic treated groups, consistent with marked progress of macroscopic wound healing processes **(Figure 8A and Figure 5)**. For a convenient understanding and visualisation of modulations in NF-κB expression, all experimental groups have been showcased in the form of a heat map based on expression intensities obtained from immunohistochemical analysis with the aid of ImageJ software. Higher, intermediate and lower expressional intensities are marked in the shades of red, orange and green, respectively **(Figure 8D)**.

**Figure 8:**
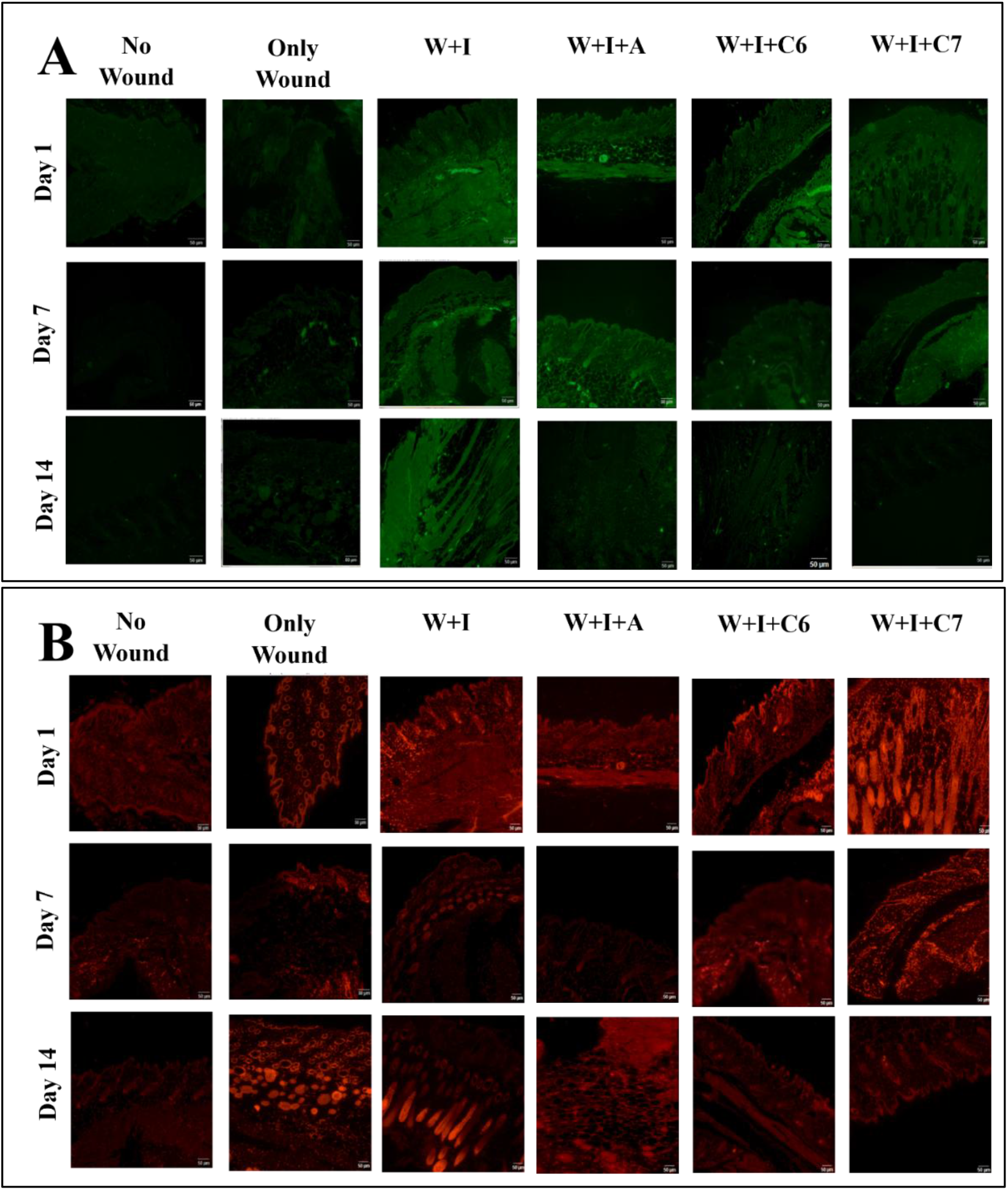
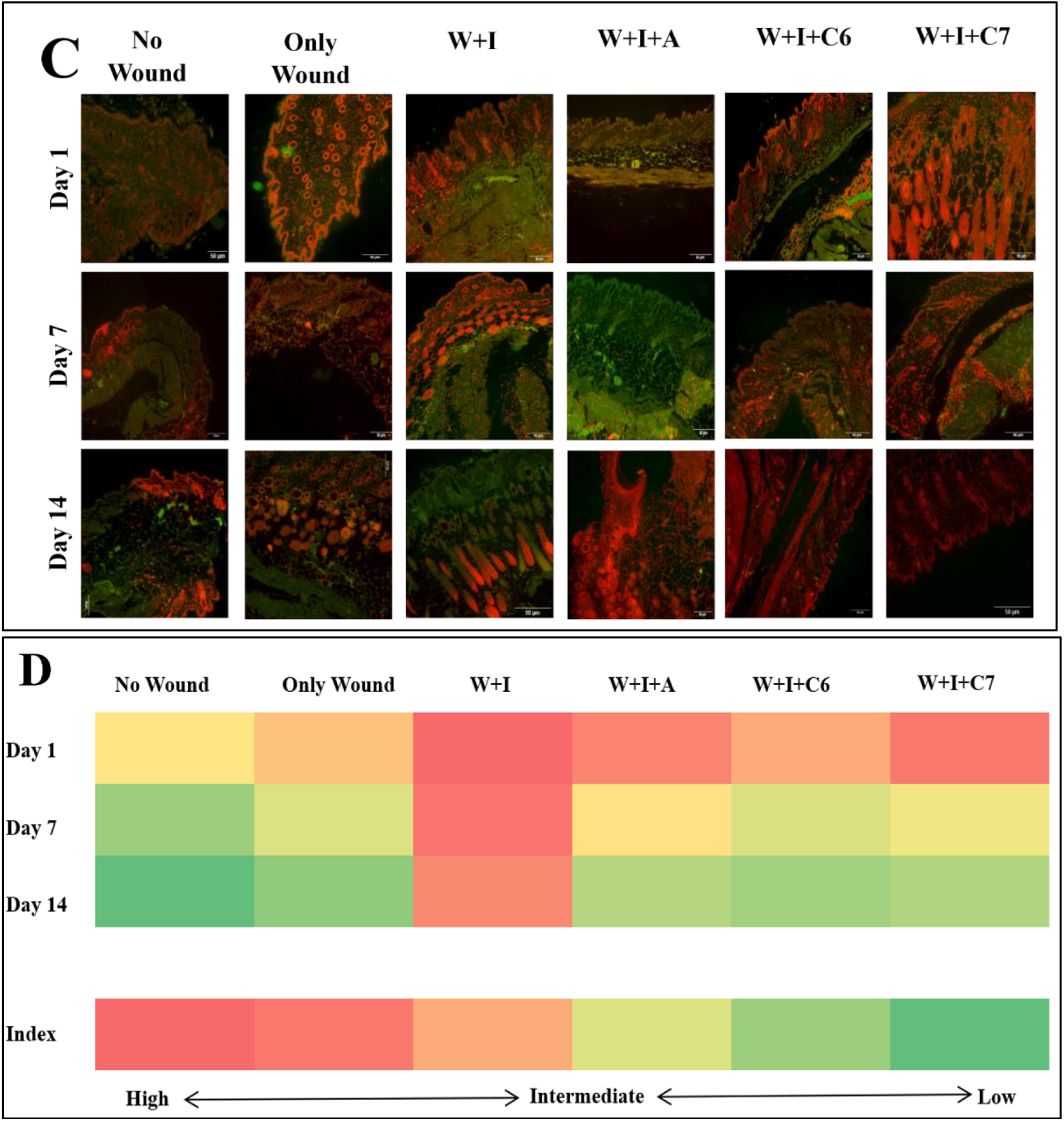
Immunohistochemical analysis of (A) NF-κB expression, (B) cellular damage assessment by propidium iodide (PI) staining and (C) merged NF-κB and PI image across experimental groups at day 1, 7, and 14. (D) Heatmap reflecting expressional intensity modulation of NF-κB protein all along the treatment period. Scale bar 50 μm.

Propidium iodide (PI) staining serves as an indirect measure of assessing cellular damage. PI can only permeate into the cell through damaged cell membrane, binds DNA and emits fluorescence when observed through fluorescence microscope. Higher PI intensity reflects more cellular damage while lower intensity reflects lesser damage, as well as healing of cellular damage. The obtained data clearly demonstrated that PI fluorescence was intensified on day 1 in all experimental groups but decreased gradually over subsequent days and minimized on day 14, corroborating well with visual wound healing characteristics. PI intensity in the C6 and C7 treated groups was less than W+I and antibiotic treated groups, suggesting that both the compounds reduced inflammation and enhanced cytoprotection **(Figure 8B)**.

Combining the NF-κB immunostaining with PI staining **(Figure 8C)**, showed that elevated expression of NF-κB was predominantly localized in PI-positive, damaged cells, indicating an association between inflammatory signaling and cellular injury. Compound C6 and C7 exhibited attenuation both in inflammation as well as cellular injury. These compounds not only impair bacterial virulence but also mitigate host bystander damage which suggests their possible role as promising antivirulence therapeutics for *S. aureus* wound infections.

## Conclusion

This study presents a comprehensive investigation of SaGpx’s plausible role in Staphylococcal virulence. For this purpose, small molecule-based inhibitors (α, β-unsaturated ketoesters) targeting SaGpx were rationally designed and synthesized. These inhibitors showed their capacity to bind SaGpx reflected by their respective K_d_ values as low as 32 µM and potent SaGpx inhibition via the FOX assay. These inhibitors demonstrated strong in vitro antibacterial activity against *S. aureus* with MIC values as low as 6.25 µM. In vivo antibacterial activity of these inhibitors was also assessed using *S. aureus* infected skin-wound mouse model where these inhibitors showed antibacterial as well as wound healing efficacy. By designing inhibitors targeting SaGpx, a possible chemical inhibition state of SaGpx was created, which may mimic the SaGpx-deficient *S. aureus* strain. However, we could not make SaGpx knock out variant, study of which would be more relevant to probe its role in *S. aureus* pathogenesis and would minimize the possibilities that the antibacterial effect of the synthesized inhibitors may also partly stem from the inhibition of other thiolate containing defense enzymes (such as peroxiredoxins) of *S. aureus* important for the bacterial pathogenesis. Future work generating an actual SaGpx knockout strain for direct comparison with the results obtained from this inhibitor-based study will further strengthen these findings and will solidify SaGpx’s antivirulence target status. Also, co-crystal structures of these compounds with SaGpx will provide insights about binding mechanism which will guide further lead optimization.

## Materials and Methods

### 1. Purification, Crystallization and Structure Solution

The SaGpx ORF (Uniprot Id: Q2FYZ0) from *Staphylococcus aureus* RN4220 strain was cloned into pET28 plasmid vector and over-expressed in E. coli BL21 (DE3). Cells grown in LB at 37º C to an O.D_600_ nm of ∼0.6 were induced with 100 µM of Isopropyl β-D-1-thiogalactopyranoside (IPTG) for 4 h at 37º C, harvested and lysed by sonication in lysis buffer (10 mM Tris pH 8.0, 300 mM NaCl and 10 mM Imidazole). The supernatant fraction was then purified via Ni-NTA affinity chromatography with different gradients of Imidazole (10 mM to 300 mM) in elution buffer and the eluted fractions containing the target protein were identified by running 15% SDS-PAGE. The selected fraction is then further purified by Superdex 75 pg size exclusion chromatography column (Cytiva) with gel filtration buffer (20 mM Tris pH 8.0, 150 mM NaCl and 5 mM DTT. Monodisperse fractions at λ_Max_. = 280 nm **(Figure S1A)** was stored at -80º C with final 5% glycerol for further assays. Site-directed mutant of SaGpx (C36S) was generated using QuikChange II Site-Directed Mutagenesis Kit (Agilent) as per the given protocol and purified identically as wild type SaGpx **(Figure S1B)**.

For Crystallization, homogenously purified SaGpx C36S mutant (22mg/mL) in 20 mM Tris pH 8.0, 250 mM NaCl and 5 mM DTT was screened via sitting-drop vapour diffusion method with Crystal Screen and Index Screen crystallization sparse matrix solutions (Hampton Research). C36S mutant crystals optimized from 2.2 M ammonium sulfate with 0.1 M Tris pH 7.0 in hanging-drop vapour diffusion method (Plate-like morphology; **Figure S2A**) grew at 298 K within 48-96 h.

Single crystals were diffracted to high resolution **(Figure S2B)** at PX-BL21 beamline facility at Indus-2, RRCAT, Indore, India ^21^. A total of 180 frames were collected with 1º oscillation by keeping the detector at 150 nm distance with continuous flow of liquid nitrogen stream at 100K temperature. Data were reduced, indexed and integrated using XDS software package ^22^. Subsequently the point group was determined, and scaling of the reduced dataset was performed by using Pointless ^23^ and SCALA ^23^ respectively from CCP4 suite ^24^. The asymmetric unit occupancy was estimated via Matthews coefficient^25^. Phasing was achieved by Molecular replacement using Phaser MR or MOLREP ^26^ of CCP4 suite with glutathione-dependent phospholipid peroxidase Hyr1 from the yeast Saccharomyces cerevisiae (PDB Id: 3CMI) ^27^ as the template. Post-phasing model underwent rigid body refinement, restrain body refinement using Refmac5 ^28^ of CCP4 suite or Phenix ^29^ and iterative manual adjustment/visualization in Coot ^30^ until data-model agreement converged. The stereochemically validated final structure was deposited to the PDB ^31^ (PDB ID: 9XP8).

### 2. Binding Affinity Analysis of Ketoester Derivatives Against SaGpx

The ketoester derivatives were designed to interact with the SaGpx’s nucleophilic active site thiol group (C36) which will subsequently render the enzyme catalytically inactive. To confirm the desired thiol modification by these compounds, a colorimetric assay employing Ellman’s reagent [5,5’-dithio-bis-(2-nitrobenzoic acid), also known as DTNB] was performed ^32^. SaGpx was buffer exchanged with reaction buffer (20 mM Tris, pH 8.0 and 150 mM NaCl) to remove residual DTT. SaGpx (20 µM) was incubated with varying concentrations (0 to 250 µM in ≤ 1% DMSO) of the synthesized ketoester compounds for 2 minutes in 250 µL total reaction volume. The reaction was stopped by adding 250 µL 10% SDS. Then 25 µL of 10 mM DTNB (prepared in DMSO) solution was added and incubated for another 30 minutes. Finally, absorbance at 412 nm was recorded in SHIMADZU UV-1900i UV-Vis Spectrophotometer. All the experiments were performed in triplicate. Mean values were plotted with standard deviation and K_d_ values were calculated using Origin software for each of the ketoester compounds tested against SaGpx.

### 3. Inhibitory Activity of Ketoester Derivatives Against SaGpx

The inhibitory activity of the ketoester derivatives against SaGpx was evaluated using FOX assay with some modifications ^33^. SaGpx was buffer exchanged with reaction buffer (20 mM Tris, pH 8.0 and 150 mM NaCl) to remove residual DTT. SaGpx (10 µM) was incubated with 100 µM t-BOOH for varying time (0 to 60 minutes) in 50 µL reaction volume. DTT (1 mM) was used as reducing agent to allow multiple turnovers. The reaction was stopped at specific timepoint by adding 950 µL of FOX working reagent and incubated for another 40 minutes. The absorbance at 560 nm was recorded in SHIMADZU UV-1900i UV-Vis Spectrophotometer. All the experiments were performed in triplicate. Mean values were plotted with standard deviation using Origin software and results were analyzed.

### 4. Antibacterial Activity of Ketoester Derivatives

#### 4.1. Agar Well Diffusion Assay

Antibacterial activity of ketoester derivatives was evaluated against *Staphylococcus aureus* MSSA476 strain. Overnight culture was subcultured (with 1% inoculum) to mid-log phase and adjusted to achieve a final working inoculum density of ∼10^5^ CFU/mL by measuring absorbance at 600 nm. LB agar plates were swabbed uniformly with the bacterial inoculum, and wells (∼ 8mm diameter) were punched into the agar using a sterile cork borer. To each well, 20 µL of the compounds (1 mM dissolved in 100% DMSO) along with Positive control (1 mM kanamycin) and negative control (100% DMSO alone) were added on each plate. This experiment was performed in triplicates for each of the compounds and controls. The inoculated plates were incubated at 37° C for 16 to 20 h in an aerobic atmosphere. After incubation, the diameter of clear inhibition zones surrounding each well was measured in centimeters using a ruler. Zone measurements were recorded from each replicate, and the data were expressed as mean inhibition zone diameter with standard deviation.

#### 4.2. Minimum Inhibitory Concentration (MIC) Assay

Following agar well diffusion screening, MICs were determined for selected ketoester derivatives by following standard broth microdilution protocol ^34^. Overnight culture *S. aureus* MSSA476 strain was subcultured (with 1% inoculum) to mid-log phase and adjusted to achieve a final working inoculum density of ∼10^5^ CFU/mL by measuring absorbance at 600 nm. Working concentration (200-1000 µM) of the compounds were prepared aseptically and freshly in DMSO and tested in 96-well plate alongside sterility (broth only), growth (inoculum only), and vehicle (broth with solvent) controls. Plates were incubated statically at 37° C for 16-20 h in an aerobic atmosphere. Following incubation, bacterial growth was assessed by taking the absorbance at 600 nm. The MIC value was defined as the lowest concentration of test compounds at which no bacterial growth was observed (i.e., well remained clear or comparable to the sterility control). All assays were performed in triplicate and MIC values were expressed as the mean of replicates.

#### 4.3. Cell Migration and Scratch Closure Assay

L929 mouse fibroblast cell line was procured from the National Centre for Cell Science (NCCS), Pune. These cells were cultured in DMEM media supplemented with 10% FBS at 37° C in a humidified environment containing 5% CO_2_. L929 cells after attaining about 95% confluency scratched using sterile 200 µL pipette tips as per standard protocol ^35^. Detached cells were removed by washing with PBS and were replenished with fresh culture media containing vehicle control (only PBS) and compound C6 and C7 (100 µM each). Plates were incubated at 37° C and images of the plates were captured at 0 h and subsequently at specified time intervals (such as 0, 2, 4, and 24 h) using an inverted phase-contrast microscope. Scratch area was analyzed via ImageJ Software, processed and plotted onto graph. Scratch closure rate (%) was calculated using the formula:@

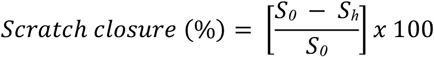

Where, S_0_ = scratch area immediately after scratching; S_h_ = Scratch area in post-scratching hours. Experiments were performed in triplicates and data are represented as mean with standard deviation.

#### 4.4. Wound Healing Efficacy Against *S. aureus* Infected Skin-wound Mice Model

##### 4.4.1. Animals

Healthy inbred strains of Swiss Albino Mice (Mus musculus), approximately 6-8 weeks old and with around 25 g weight, were chosen for this experiment. All the experimental animals were acclimatized for 15 days in an environmentally conditioned room where 25±2°C temperature, 55±5% humidity and a 12-hour light/dark cycle were maintained with unimpeded access to food and water. Animal handling during the experiment was carried out under the guidance and supervision of the Committee for the Control and Supervision of Experiments on Animals (CPCSEA newly named as CCSEA). Institutional Animal Ethical Committee, University of Kalyani, West Bengal, India (IACE-KU) approved protocols were followed in this experiment (approval number: 892/GO/Re/S/01/CPCSEA and 892/GO/Re/S/2005/CCSEA).

##### 4.4.2. Experimental Design

For this experiment, six different murine groups were taken as described below:

i. Group 1: No wound - Mice group with no wounds.
ii. Group 2: Only Wound - An Excisional wound was generated on the back of healthy mice and no inhibitors applied.
iii. Group 3: Wound + Infection (W+I) - Mice group with excisional wound infected with *S. aureus* monitored without any inhibitor application.
iv. Group 4: Wound + Infection + Antibiotic (W+I+A): In this group of mice, a known antibiotic (Neomycin) was applied to the *S. aureus*-infected wound.
v. Group 5: Wound + Infection+ Compound C6 (W+I+C6): C6 was applied on *S. aureus* infected excisional wound.
vi. Group 6: Wound + Infection + Compound C7 (W+I+C7): C7 was applied on *S. aureus*-infected excisional wound.

##### 4.4.3. Excisional Wounding of Experimental Animals

After shaving the fur on the dorsal surface of mice, an excisional wound of approximately 8 mm in diameter was created using a sterile pair of scissors and forceps. The wound area was wiped off with Isopropyl alcohol. Mice were anaesthetized during the procedure to reduce suffering.

##### 4.4.4. Loading of Bacteria in Excisional Wound

Immediately after wounding, 50 µL bacterial suspension adjusted to ∼10^8^ CFU/mL was carefully applied onto the wound, which was then covered with sterile gauze for the development of infection. Animals were observed for the next 24 h for the development of visible symptoms of infection.

##### 4.4.5. Inhibitor Dose Selection and Application

A range-finding trial with inhibitor concentrations of 75 µM, 85 µM and 100 µM was carried out by using cytotoxicity assay following standard protocol ^36^. Among different concentrations, 100 µM inhibitor concentration was found to be effective. After the appearance of visual infection symptoms on the wounded mice, 100 µL of inhibitor (100 µM concentration) was applied in accordance with the experimental design and monitored thereafter at regular interval.

##### 4.4.6. Measurement of Wound Area and Percentage of Wound Area Contraction

Wound edges were traced onto sterilized transparent sheet cautiously on the day of excision, day after inoculation, inhibitor application day 1, 7 and 14 respectively and measured using mm^2^ graph paper. Digital photographs were clicked on the aforementioned days to track the changes in overall wound visually. Wound area contraction rate (%) was calculated by the following formula:

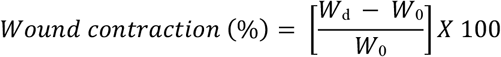

Where, W_d_ = wound area in post-wounding day; W_0_ = wound area in the wounding day.

##### 4.4.7. Histological Preparations

Mice from each experimental group was euthanized as per standard protocol ^37^. Skin of wounded area were taken into 10% neutral buffered formalin directly and fixed for 48 h, dehydrated and embedded in paraffin. Paraffin embedded skin tissue was cut into 5 micrometer sections and subjected to Hematoxylin-Eosin staining and immunohistochemical procedures ^38,39^.

##### 4.4.8. Immunohistochemistry

Expression levels of pro-inflammatory marker protein were estimated using FITC(Fluorescein-5-isothiocyanate) tagged secondary anti-mouse antibody (procured from Santa Cruz Biotechnology, USA). Immunofluorescent photographs were acquired using Carl Zeiss Axio Vert A1 microscope and analyzed through Image J software ^38–40^.

##### 4.4.9. Assessment of DNA Damage by Propidium Iodide (PI) Staining

Paraffinized histological skin tissue sections were deparaffinized, rehydrated, incubated with primary and FITC (Fluorescein-5-isothiocyanate) tagged secondary antibody sequentially, followed by counterstaining with Propidium Iodide.

##### 4.4.10. Statistical Analysis

Statistical significance of results obtained from triplicated experiments were assessed through Student’s t-test after calculation of mean and standard error. ***p<0.001,**p<0.01, *p<0.05 were considered statistically significant for this experiment.

## Acknowledgements

The study is supported by grants from Start-up Research Grant (SRG) funded by Science and Engineering Research Board (SERB), Department of Science and Technology, Government of India (Sanction no. SRG/2020/001353), and AYURTECH facility at IIT Jodhpur, Ministry of AYUSH, Government of India (Sanction no. S-12011/12/2021-SCHEME). S.M. expresses special thanks to Ministry of Education, Government of India for providing research fellowships to carry out this work. Authors are thankful to IIT Jodhpur for providing experimental facility and necessary infrastructure for carrying out this work. Authors are also thankful to PX-BL21 beamline (BARC) at Indus-2, RRCAT, Indore, India for protein crystal diffraction and subsequent data collection facility. Authors sincerely thank S.N Bose Innovation Centre, University of Kalyani for providing central instrumentation facility used for data curation.

## Author Contributions

**SM:** Investigation, Conceptualization, Methodology, Data curation, Formal analysis, Writing – original draft. **SD:** Investigation, Methodology, Data curation, Formal analysis, Writing – original draft. **AK:** Investigation, Methodology, Data curation, Formal analysis, Writing – original draft. **HS:** Methodology, Data curation **NS:** Visualization, Formal analysis. **AS:** Supervision, Investigation, Resources, Writing – review and editing. **NKR:** Supervision, Investigation, Resources, Writing – review and editing. **SB:** Conceptualization, Project administration, Supervision, Writing – review and editing, Resources and Funding acquisition.

