## Supplementary Data for "Exploring the Structural and Functional role of α,β-unsaturated Ketoesters as Anti-Staphylococcal Agents Targeting Glutathione Peroxidase"

Sushobhan Maji[1], Sayoni Dam[2], Akanksha Kumari[3], Himani Sharma[3], Naveen Sharma[4], Nirmal K. Rana[3*], Asmita Samadder[2], Sudipta Bhattacharyya[1*]
[1]Department of Bioscience and Bioengineering, Indian Institute of Technology Jodhpur, Jodhpur 342037, India
[2]Department of Zoology, University of Kalyani, Kalyani 741235, India
[3]Department of Chemistry, Indian Institute of Technology Jodhpur, Jodhpur 342037, India
[4]Discovery Research Division, Indian Council of Medical Research, New Delhi 110029, India

[*]Correspondence to:
Sudipta Bhattacharyya, Department of Bioscience & Bioengineering, Indian Institute of Technology, Jodhpur 342030, Rajasthan, India.
Nirmal K. Rana, Department of Chemistry, Indian Institute of Technology, Jodhpur 342030, Rajasthan, India.

### 2. Materials and Methods:

#### 2.1. Fine Chemicals, Reagents and Enzymes:

All the fine chemicals and reagents needed for biological evaluation of synthesized small molecule-based inhibitors described in this study were of analytical grades and procured from SRL (India), Sigma-Aldrich (USA) and other chemical vendors with international repute. Antibodies used in Immunohistochemical studies were purchased from Santa Cruz Biotechnology, USA.

#### 2.2. General Synthetic Procedure of Ketoester derivatives:[1]

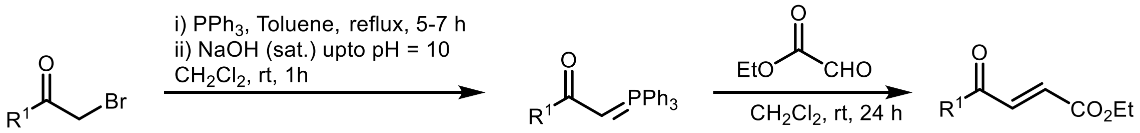


A solution of α-bromoketone (5 mmol, 1.0 equiv.) in toluene (40 mL) was treated with triphenylphosphine (1.31 g, 5 mmol, 1.0 equiv.), and the reaction mixture was heated under reflux for 6 h. After completion, the resulting solid was collected by filtration and dissolved in dichloromethane. A saturated aqueous sodium hydroxide solution was then added until the pH reached ~10, and the mixture was stirred at room temperature for 1 h. The organic layer was extracted with dichloromethane (3 × 70 mL), and the combined extracts were washed with brine (3 × 100 mL), dried over anhydrous sodium sulfate, filtered, and concentrated under reduced pressure to afford a crude product, which was used directly in the next step without further purification.
In a separate flask, ethyl glyoxylate (5 mmol, 1.0 equiv.) was dissolved in dichloromethane (40 mL), and the above crude intermediate was added at room temperature. The reaction mixture was stirred for 24 h, after which the solvent was removed under reduced pressure. The residue was purified by flash column chromatography on silica gel (100–200 mesh) using hexane/ethyl acetate (19:1) as the eluent to afford the desired product.

**Characterization of α,β-unsaturated-β-keto esters.**

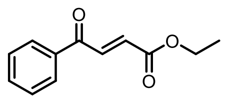

**Ethyl-4-oxo-4-phenylbut-2-enoate (C1)**[1] **:** $^1$H NMR (500 MHz, $CDCl_3$) δ 7.97 – 7.90 (m, 2H), 7.84 (d, *J* = 15.6 Hz, 1H), 7.56 (t, *J* = 7.4 Hz, 1H), 7.45 (t, *J* = 7.7 Hz, 2H), 6.81 (d, *J* = 15.6 Hz, 1H), 4.24 (q, *J* = 7.1 Hz, 2H), 1.28 (t, *J* = 7.1 Hz, 3H); HRMS (ESI) m/z: [M + H]+ calcd for $C_{12}H_{13}O_3$ 205.0860; found: 205.0896.

**Ethyl-4-(4-fluorophenyl)-4-oxobut-2-enoate (C2)**[1] **:**

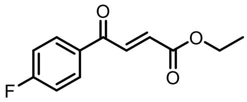

$^1$H NMR (500 MHz, $CDCl_3$) δ 8.08 – 7.95 (m, 2H), 7.85 (d, *J* = 15.5 Hz, 1H), 7.16 (t, *J* = 8.6 Hz, 2H), 6.85 (d, *J* = 15.5 Hz, 1H), 4.28 (q, *J* = 7.1 Hz, 2H), 1.32 (t, *J* = 7.1 Hz, 3H); HRMS (ESI) m/z: [M + H]+ calcd for $C_{12}H_{12}FO_3$ 223.0765; found: 223.0771.

**Ethyl-4-(4-bromophenyl)-4-oxobut-2-enoate (C3)**[1] **:**

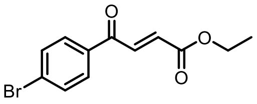

$^1$H NMR (500 MHz, $CDCl_3$) δ 7.89 – 7.81 (m, 3H), 7.69 – 7.63 (m, 2H), 6.89 (d, *J* = 15.5 Hz, 1H), 4.30 (q, *J* = 7.1 Hz, 2H), 1.35 (t, *J* = 7.1 Hz, 3H); HRMS (ESI) m/z: [M + H]+ calcd for $C_{12}H_{12}BrO_3$ 282.9965 & 284.9944; found: 284.9961.

**Ethyl-4-(4-cyanophenyl)-4-oxobut-2-enoate (C4)**[1] **:**

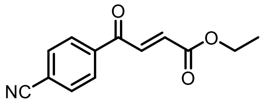

$^1$H NMR (500 MHz, $CDCl_3$) δ 8.07 (d, *J* = 8.1 Hz, 2H), 7.83 (t, *J* = 10.5 Hz, 3H), 6.90 (d, *J* = 15.5 Hz, 1H), 4.30 (q, *J* = 7.1 Hz, 2H), 1.34 (t, *J* = 7.1 Hz, 3H); HRMS (ESI) m/z: [M + H]+ calcd for $C_{13}H_{12}NO_3$ 230.0812; found: 230.0834.

**Ethyl-4-(3,5-difluorophenyl)-4-oxobut-2-enoate (C5)**[1] **:**

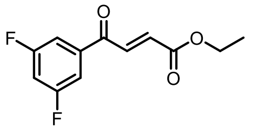

$^1$H NMR (500 MHz, $CDCl_3$) δ 7.89 (td, *J* = 8.5, 6.5 Hz, 1H), 7.72 (dd, *J* = 15.5, 3.5 Hz, 1H), 7.05 – 6.96 (m, 1H), 6.91 (ddd, *J* = 11.0, 8.6, 2.4 Hz, 1H), 6.83 (dd, *J* = 15.5, 1.3 Hz, 1H), 4.28 (q, *J* = 7.1 Hz, 2H), 1.33 (t, *J* = 7.1 Hz, 3H); HRMS (ESI) m/z: [M + H]+ calcd for $C_{12}H_{11}F_2O_3$ 241.0671; found: 241.0682

**Ethyl-4-(2,4-dichlorophenyl)-4-oxobut-2-enoate (C6)**[1] **:**

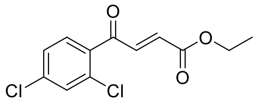

$^1$H NMR (500 MHz, $CDCl_3$) δ 7.51 (d, *J* = 15.8 Hz, 1H), 7.47 (dd, *J* = 6.5, 5.2 Hz, 2H), 7.35 (dd, *J* = 8.3, 1.9 Hz, 1H), 6.66 (d, *J* = 15.8 Hz, 1H), 4.28 (q, *J* = 7.1 Hz, 2H), 1.32 (t, *J* = 7.1 Hz, 3H); HRMS (ESI) m/z: [M + H]+ calcd for $C_{12}H_{11}Cl_2O_3$ 273.0080; found: 273.0095

**Ethyl-4-(3,5-dichlorophenyl)-4-oxobut-2-enoate (C7)**[1] **:**

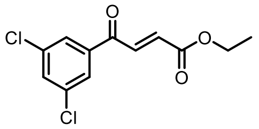

$^1$H NMR (500 MHz, $CDCl_3$) δ 7.70 (d, *J* = 16.1 Hz, 1H), 7.49 (d, *J* = 1.8 Hz, 2H), 7.42 (t, *J* = 1.8 Hz, 1H), 7.36 (d, *J* = 16.1 Hz, 1H), 4.40 (q, *J* = 7.1 Hz, 2H), 1.42 (t, *J* = 7.2 Hz, 3H); HRMS (ESI) m/z: [M + H]+ calcd for $C_{12}H_{11}Cl_2O_3$ 273.0080; found: 273.0091

**Ethyl-4-(3,4-dichlorophenyl)-4-oxobut-2-enoate (C8)**[1] **:**

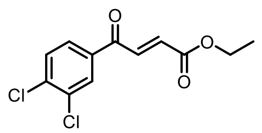

$^1$H NMR (500 MHz, $CDCl_3$) δ 8.08 (d, *J* = 2.0 Hz, 1H), 7.86 – 7.78 (m, 2H), 7.60 (d, *J* = 8.4 Hz, 1H), 6.91 (d, *J* = 15.5 Hz, 1H), 4.31 (q, *J* = 7.1 Hz, 2H), 1.36 (t, *J* = 7.1 Hz, 3H); HRMS (ESI) m/z: [M + H]+ calcd for $C_{12}H_{11}Cl_2O_3$ 273.0080; found: 273.0102

**Ethyl-4-([1,1'-biphenyl]-4-yl)-4-oxobut-2-enoate (C9)[1] :**

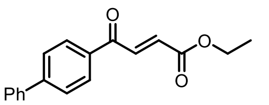

$^1$H NMR (500 MHz, $CDCl_3$) δ 8.11 (d, *J* = 8.4 Hz, 2H), 7.98 (d, *J* = 15.5 Hz, 1H), 7.76 (d, *J* = 8.4 Hz, 2H), 7.70 – 7.64 (m, 2H), 7.51 (t, *J* = 7.5 Hz, 2H), 7.44 (t, *J* = 7.3 Hz, 1H), 6.95 (d, *J* = 15.5 Hz, 1H), 4.34 (q, *J* = 7.1 Hz, 2H), 1.39 (t, *J* = 7.1 Hz, 3H); HRMS (ESI) m/z: [M + H]+ calcd for $C_{18}H_{17}O_3$ 281.1173; found: 281.1205

**Ethyl-4-(naphthalen-2-yl)-4-oxobut-2-enoate (C10)[1] :**

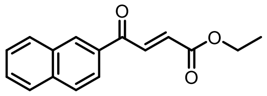

$^1$H NMR (500 MHz, $CDCl_3$) δ 8.46 (s, 1H), 8.02 (t, *J* = 11.0 Hz, 2H), 7.98 – 7.79 (m, 3H), 7.55 (dt, *J* = 14.9, 6.9 Hz, 2H), 6.91 (d, *J* = 15.5 Hz, 1H), 4.29 (dd, *J* = 14.2, 7.1 Hz, 2H), 1.33 (t, *J* = 7.1 Hz, 3H); HRMS (ESI) m/z: [M + H]+ calcd for $C_{16}H_{15}O_3$ 255.1016; found: 255.1049

**Ethyl-5,5-dimethyl-4-oxohex-2-enoate (C11)[1] :**

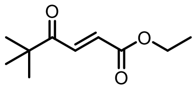

$^1$H NMR (500 MHz, $CDCl_3$) δ 7.49 (d, *J* = 15.5 Hz, 1H), 6.75 (d, *J* = 15.4 Hz, 1H), 4.24 (q, *J* = 7.1 Hz, 2H), 1.30 (t, *J* = 7.1 Hz, 3H), 1.17 (s, 9H); HRMS (ESI) m/z: [M + H]+ calcd for $C_{10}H_{17}O_3$ 185.1173; found: 185.1164

**Ethyl-4-((3r,5r,7r)-adamantan-1-yl)-4-oxobut-2-enoate (C12)[1] :**

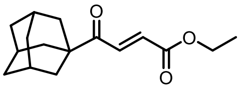

$^1$H NMR (500 MHz, $CDCl_3$) δ 7.53 (d, *J* = 15.5 Hz, 1H), 6.74 (d, *J* = 15.5 Hz, 1H), 4.26 (q, *J* = 7.1 Hz, 2H), 2.08 (s, 3H), 1.83 (d, *J* = 2.4 Hz, 5H), 1.74 (dd, *J* = 34.8, 12.1 Hz, 7H), 1.32 (t, *J* = 7.1 Hz, 3H); HRMS (ESI) m/z: [M + H]+ calcd for $C_{16}H_{23}O_3$ 263.1642; found: 263.1673

**Ethyl-4-oxopent-2-enoate (C13)[2] :**

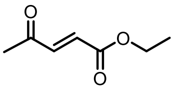

$^1$H NMR (500 MHz, $CDCl_3$) δ 6.95 (d, *J* = 16.1 Hz, 1H), 6.59 (d, *J* = 16.1 Hz, 1H), 4.21 (q, *J* = 7.1 Hz, 2H), 2.31 (s, 3H), 1.27 (t, *J* = 7.1 Hz, 3H); HRMS (ESI) m/z: [M + H]+ calcd for $C_7H_{11}O_3$ 143.0703; found: 143.0715

### 3. Results and Discussion:

**Table S1.** Data collection and refinement statistics.

| Parameters | C36S mutant of SaGpx (PDB ID: 9XP8) |
|---|---|
| **Wavelength** | 0.978930 Å |
| **Resolution range** | 39.6 - 1.65 (1.74 - 1.65) |
| **Space group** | C 2 2 21 |
| **Unit cell** | 62.76 95.73 60.33 90 90 90 |
| **Total reflections** | 158317 (23346) |

| | |
|---|---|
| **Unique reflections** | 22254 (3211) |
| **Multiplicity** | 7.1 (7.3) |
| **Completeness (%)** | 100 (100) |
| **Mean I/sigma(I)** | 20.4 (4.0) |
| **Wilson B-factor** | 17.86 |
| **R-merge** | 0.054 (0.513) |
| **R-meas** | 0.058 (0.552) |
| **R-pim** | 0.022 (0.203) |
| **CC1/2** | 0.999 (0.938) |
| **CC*** | 1 (0.983) |
| **Reflections used in refinement** | 21442 (2182) |
| **Reflections used for R-free** | 1026 (101) |
| **R-work** | 0.1815 (0.2018) |
| **R-free** | 0.2044 (0.2411) |
| **CC(work)** | 0.962 (0.951) |
| **CC(free)** | 0.965 (0.884) |
| **Number of non-hydrogen atoms** | 1504 |
| **macromolecules** | 1279 |
| **ligands** | 44 |
| **solvent** | 193 |
| **Protein residues** | 158 |
| **RMS(bonds)** | 0.012 |
| **RMS(angles)** | 1.15 |
| **Ramachandran favored (%)** | 98.08 |
| **Ramachandran allowed (%)** | 1.92 |
| **Ramachandran outliers (%)** | 0.00 |
| **Rotamer outliers (%)** | 0.00 |
| **Clashscore** | 4.67 |
| **Average B-factor** | 27.62 |
| **macromolecules** | 25.34 |
| **ligands** | 65.09 |
| **solvent** | 36.56 |
| **Number of TLS groups** | 1 |

Statistics for the highest-resolution shell are shown in parentheses.

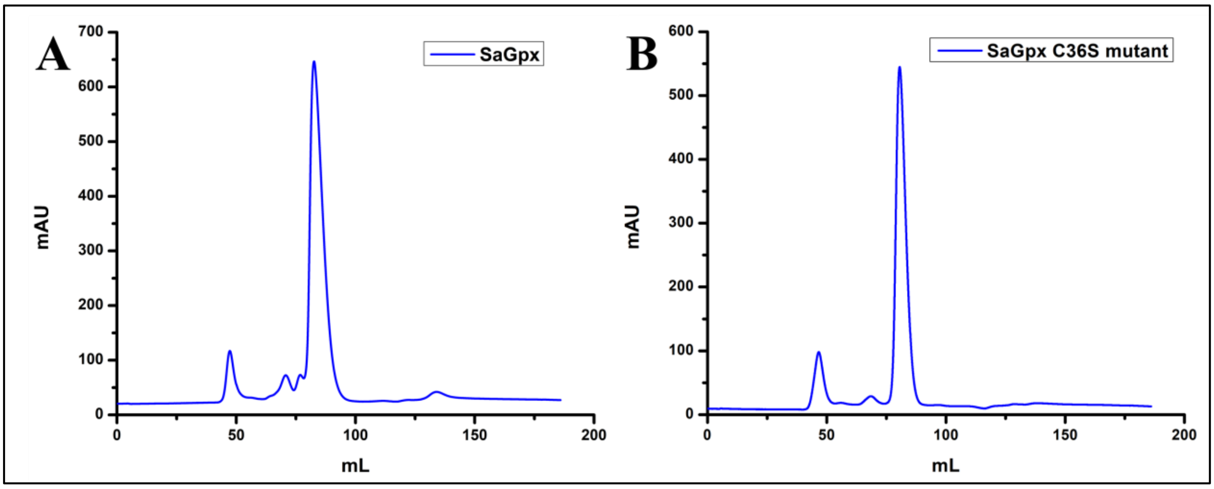


**Figure S1:** Size-exclusion chromatography of purified SaGpx and SaGpx C36S mutant yielded a single homogenous peak at $\lambda_{Max}$ = 280nm, corresponding to the target protein fraction. Thess monodisperse protein sample was collected for subsequent biochemical characterization as well as crystallization purposes.

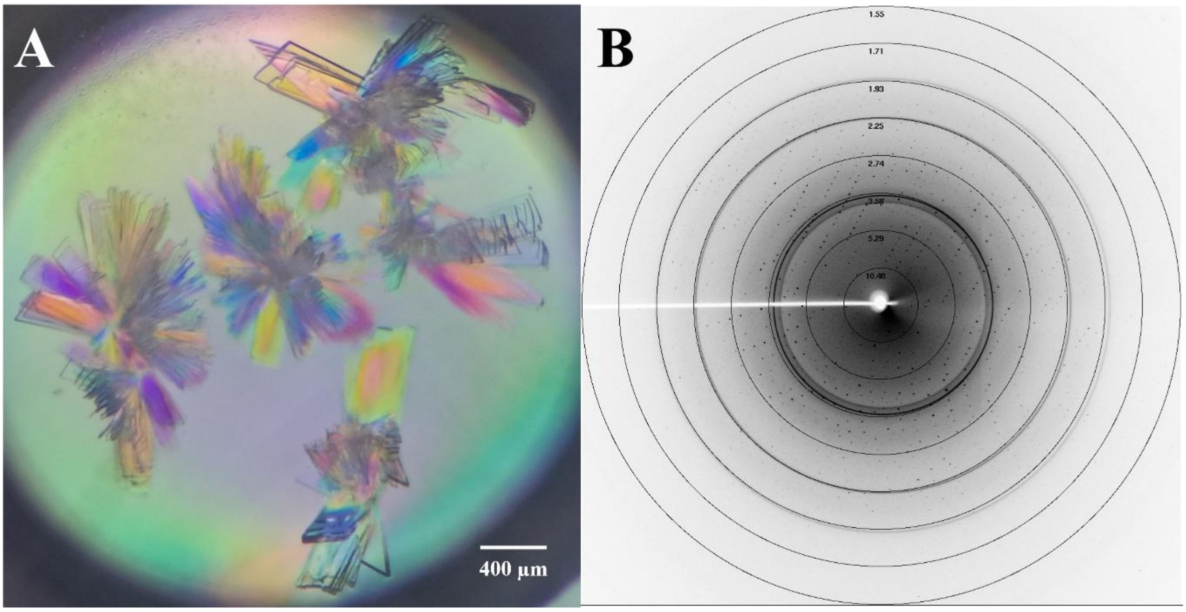


**Figure S2:** Crystals and single-crystal x-ray diffraction of SaGpx (C36S mutant). (A) Crystal drop shows plate like crystals radiating from a hub. Crystals obtained from hanging drop vapour diffusion method method. (B) X-ray diffraction pattern of single crystal. The well-defined sharp diffraction spots with low background noise indicate high crystal quality, with diffraction extending upto1.5Å.

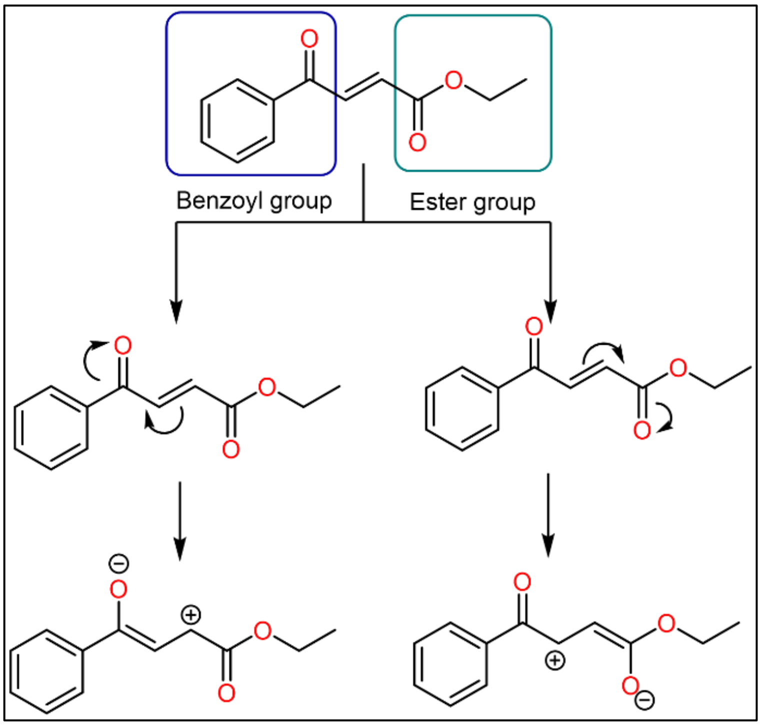


**Figure S3.** Possible mechanism of electron delocalization by the carbonyl groups present on either side of the double bond.

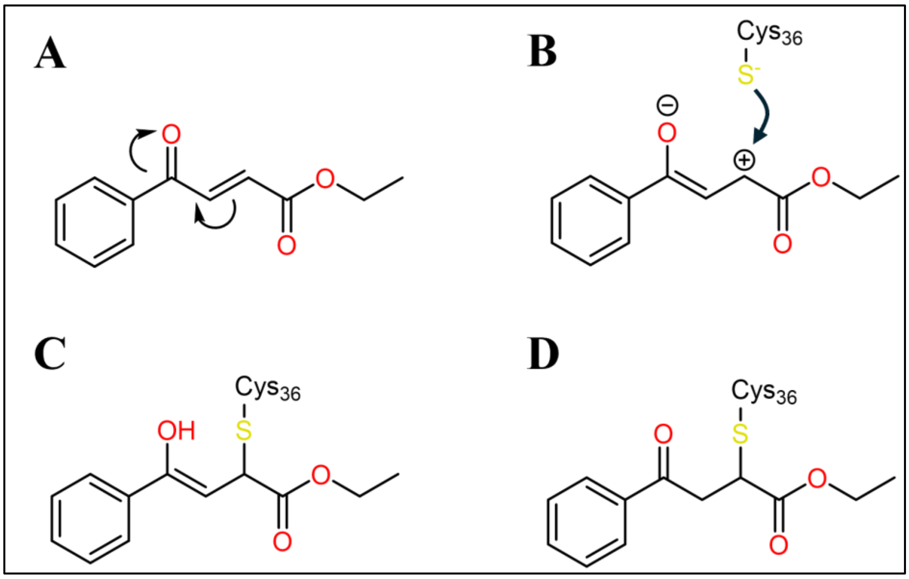


**Figure S4.** (A-D) Possible mechanism of inhibitor action with nucleophilic thiol group present in SaGpx.

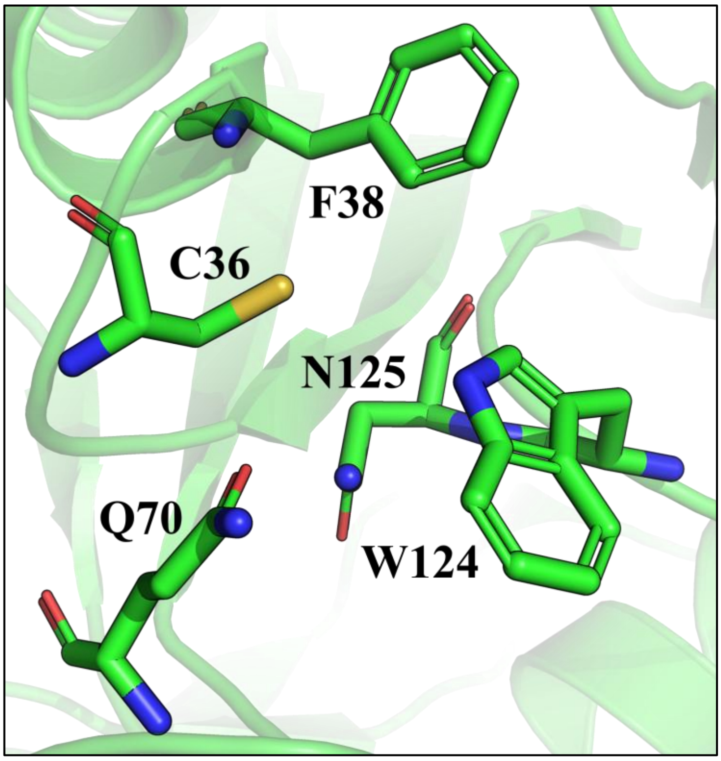


**Figure S5.** Cartoon representation of amino acid residues surrounding the C36. F38 and W124 contributes to the hydrophobic environment near the active site.

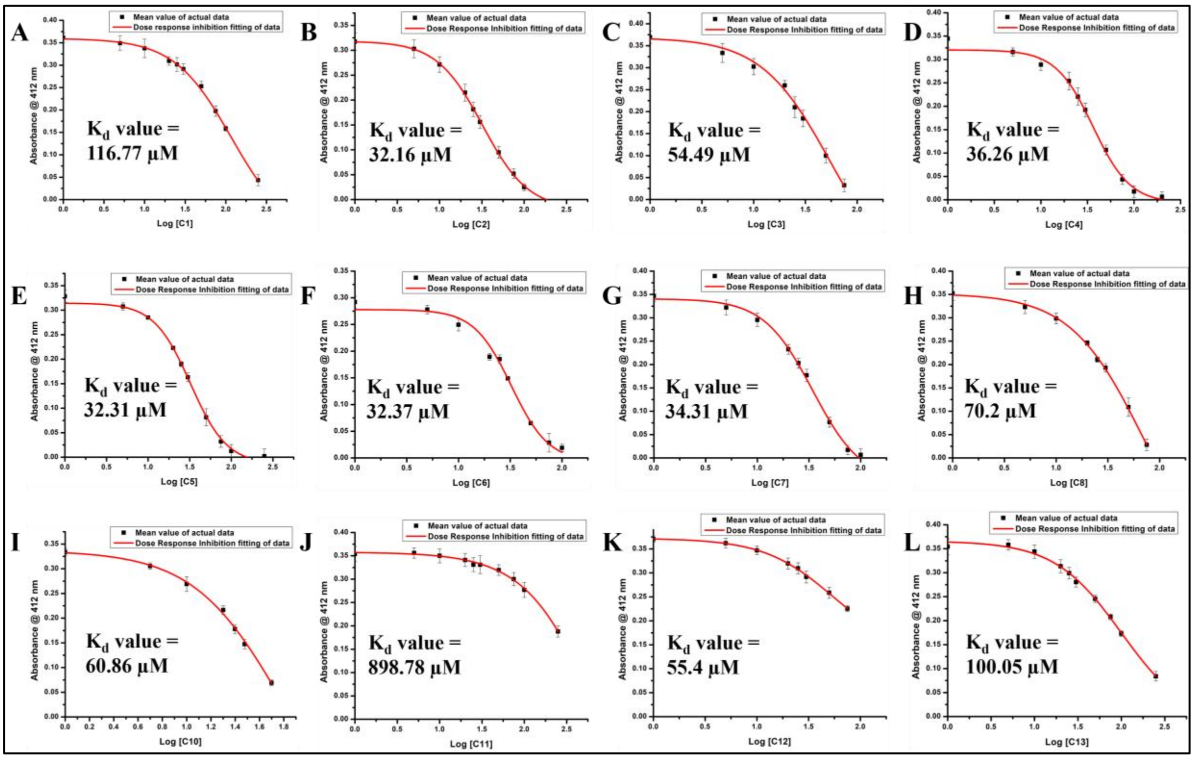


**Figure S6.** Binding affinity ($K_d$ Values) assessment of the Synthesized ketoester Derivatives Against SaGpx.

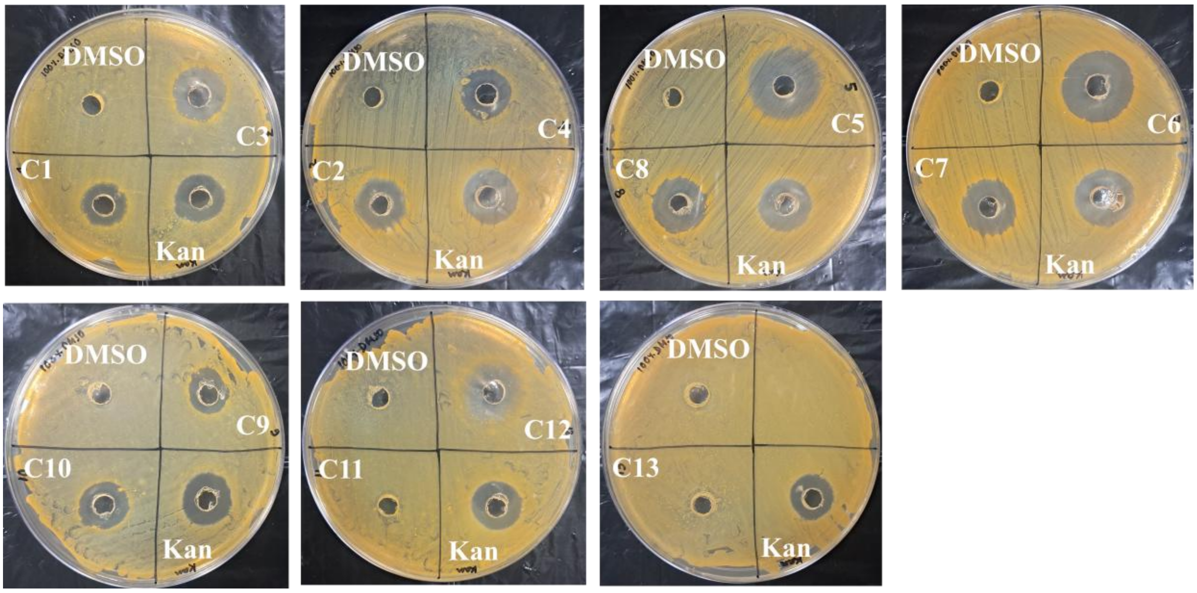


**Figure S7.** Agar well diffusion assay plates showing zones of inhibition (ZOI) against S. aureus.
